# SETD1A-dependent EME1 transcription drives PARPi sensitivity in HR deficient tumour cells

**DOI:** 10.1101/2024.06.16.599181

**Authors:** Ellie Sweatman, Rachel Bayley, Richad Selemane, Martin R. Higgs

## Abstract

**Background:** Cells deficient in DNA repair factors breast cancer susceptibility 1/2 (BRCA1/2) or ataxia-telangiectasia mutated (ATM) are sensitive to poly-ADP ribose polymerase (PARP) inhibitors. Building on our previous findings, we asked how the lysine methyltransferase SETD1A contributed to PARP inhibitor-mediated cell death and determined the mechanisms responsible.

**Methods:** We used cervical, breast, lung and ovarian cancer cells bearing mutations in *BRCA1* or *ATM* and depleted SETD1A using siRNA or CRISPR/Cas9. We assessed the effects of the PARPi Olaparib on cell viability, homologous recombination, and DNA repair. We assessed underlying transcriptional perturbations using RNAseq. We also used data from The Cancer Genomics Atlas (TCGA) to investigate overall patient survival.

**Results:** Loss of SETD1A from both BRCA1-deficient and ATM-deficient cancer cells was associated with resistance to Olaparib, explained by an partial restoration of homologous recombination. Mechanistically, SETD1A-dependent transcription of the crossover junction endonuclease EME1 correlated with sensitivity to Olaparib in these cells. Accordingly, when SETD1A or EME1 was lost, BRCA1 or ATM-mutated cells became resistant to Olaparib, and homologous recombination was partially restored.

**Conclusions:** Loss of SETD1A or EME1 may explain why patients develop resistance to PARP inhibitors in the clinic.

## 1. INTRODUCTION

Poly-ADP-ribose polymerase (PARP) inhibitors (PARPi) were first identified as potential targeted cancer treatments for tumours with deficiencies in the homologous recombination (HR) pathway of double strand break (DSB) repair^1^. This is exemplified by the exquisite sensitivity of cells bearing mutations in breast cancer susceptibility 1 (*BRCA1*) or 2 (*BRCA2*) to PARP inhibition^2^. To date, four different PARP inhibitors (Olaparib, Rucaparib, Talazoparib and Niraparib) have received clinical approval for the treatment of HR-deficient breast^3^, ovarian^4^, pancreatic^5^ and prostate cancer^6,7^. This ranges from widespread use alongside conventional chemotherapies to a more limited use in metastatic settings after failure of other treatment options.

Unfortunately, despite the success of PARPi in the clinic, patient prognosis is frequently hampered by the development of resistance. Indeed, approximately 40% of metastatic breast cancer patients harbouring germline *BRCA1/2* mutations fail to respond to Olaparib treatment^3^, suggesting widespread pre-existing resistance. This emphasises the need to further characterise PARPi resistance mechanisms, and to identify strategies to overcome resistance.

From extensive *in vitro* studies, several mechanisms of resistance to PARPi have been described ^e.g.8-10^. These include hypomorphic secondary mutations in *BRCA1/2* that partially restore function, perturbation of PARP1/2 or PARG expression, or re-wiring of the DNA damage response. In this latter case, alterations in fork protection, use of alternative error-prone pathways such as alt-EJ or the repair/prevalence of ssDNA gaps all cause PARPi resistance in BRCA-deficient cells. One of the most well-studied resistance mechanisms to PARPi is the loss of pro-NHEJ factors, which drives partial restoration of functional HR. This includes loss of 53BP1^11,12^ or inactivation of the downstream effectors RIF1, REV7 and Shieldin^13-18^. Crucially, loss of this pathway and concomitant restoration of HR has been observed in mouse models of *BRCA1* deficiency, in patient-derived xenograft models, and in patients^12,19-21^.

Recently our lab identified the lysine methyltransferase, SET domain containing protein 1 A (SETD1A) as a novel anti-resection factor that suppresses BRCA1-mediated DNA end-resection and which is vital for RIF1-dependent NHEJ^22^. Loss of SETD1A abrogates RIF1 chromatin binding and compromises NHEJ, facilitating BRCA1-independent DNA end resection in BRCA1-deficient cells. This enables the partial restoration of HR and resolution of PARPi-induced DSBs, conferring PARPi resistance *in vitro*^22,23^.

Whilst much research has focused on the impact of PARPi in BRCA-deficient cells, the clinical efficacy of PARPi extends beyond BRCA to include mutations in other DNA repair proteins involved in HR, such as the RAD51 recombinase, RAD51 paralogs C and D (RAD51C/D), excision repair cross-complementation group 1 (ERCC1), partner and localiser of BRCA2 (PALB2), and ataxia-telangiectasia mutated (ATM)^24-27^. ATM is a phosphatidyl inositol 3 kinase-like kinase that plays a key role in initiating DNA repair. ATM is a known cancer susceptibility gene that is mutated in 5% of all cancers, including 40% of mantle cell lymphomas, 20% of colorectal cancers and 10% prostate and lung cancers^28,29^. Early studies demonstrated that depletion of ATM induces PARPi sensitivity, indicating that ATM-mutated cancers could also be candidates for PARPi therapy^30^. Subsequently, *ATM*-deficient lung, gastric, prostate, brain and lymphoid tumours have all been shown to exhibit PARPi sensitivity^31-35^. However, the use of ATM as a clinical biomarker for PARPi sensitivity is still limited, and some trials in non-small cell lung cancer in which ATM is frequently mutated have proved inconclusive^36^. Moreover, whilst mechanisms of PARPi resistance are well studied in the context of BRCA-deficiency, there is a lack of understanding as to whether these apply to other HR-deficient contexts including *ATM* deficiency.

In this study we ask whether SETD1A-induced PARPi resistance is specific to BRCA1-deficiency or whether it could be broadly applicable to other HR deficient cancers, focusing on those possessing *ATM* or *BRCA2* mutations. To do so, we assessed cellular survival and DNA repair kinetics in response to Olaparib in a range of HR-deficient cell lines in the presence/absence of SETD1A. Based on its role as a lysine methyltransferase targeting H4K4, and utilising CRISPR/Cas9 to knock out *SETD1A*, we performed RNA sequencing analysis to identify the transcriptional mechanisms that underpin SETD1A-induced PARPi resistance.

Our findings expand the role of SETD1A in PARPi resistance to *ATM*-deficient cancer cells. Interestingly, SETD1A-dependent transcription of the crossover endonuclease EME1 was common in PARPi-resistant models *in vitro*, and depletion of EME1 from HR-deficient cells drove resistance to Olaparib. SETD1A and EME1 expression and/or activity could therefore offer a potential new combinatorial biomarker to help stratify patients who will benefit from PARPi therapy, reducing the heterogenous response observed in the clinic and improving patient outcomes.

## 2. MATERIALS AND METHODS

### 2.1 Cell lines and culture

All cell lines were maintained at 37 °C and 5 % CO^2^ and passaged by trypsinisation at sub-confluency. HeLa (CVCL_0030; ATCC), HeLa-H3 GFP (WT and K4A)^37,38^, MCF-7 (ATCC; CVCL_ 0031) and A549 (ATCC; CVCL_0023) cell lines were cultured in Dulbecco’s modified Eagle’s media (DMEM), supplemented with foetal calf serum (FCS) (10% v/v). H23 cell lines (ATCC; CVCL_1547) were cultured in RPMI-1640 media containing L-glutamine and supplemented with FCS (10% v/v), 1 mM sodium pyruvate and 10 mM HEPES. SKOV3 (ATCC; CVCL_0532) cell lines were cultured in McCoy 5A (Modified) Medium supplemented with FCS (10% v/v). UWB1.289 (ATCC; CVCL_B079) cell lines were cultured in RPMI-1640 media containing Mammary Epithelial Growth Medium (Lonza) (50% v/v) supplemented with FCS (3% v/v) and L-glutamine (50% v/v). HCC-1937 (DSMZ; CVCL_0290) cell lines were cultured in RPMI-1640 media containing L-glutamine supplemented with FCS (10% v/v). HeLa Kyoto iCas9^22^ cells transduced with a lentivirus expressing sgRNA targeting human SETD1A (AGAGCCATCGGAAATTTCCG) or a non-targeting control were obtained from Dr Simon Boulton. They were cultured in DMEM supplemented with tetracycline-free FCS (10% v/v). Cas9 expression was induced with 1 µg/ml doxycycline for 48 h. All media and additives were from Life Technologies unless specified.

### 2.2 siRNA transfections

SMARTpool siRNAs (Horizon Discovery) were transfected into cells using Oligofectamine (Life Technologies) at a final concentration of 100 nM following the manufacturer’s instructions. siRNA targeting LacZ (Horizon Discovery; CGUACGCGGAAUACUUCGdTdT) was used as a control.

### 2.3 Antibodies

The following antibodies were used in this study: RAD51 (Millipore; Cat# PC130), SETD1A (Bethyl; Cat# A300-289A), ATM (Santa Cruz; sc-53173), BRCA1 (Santa Cruz; Cat # sc-6954), BRCA2 (Santa Cruz; sc-6954), PARP1 (Santa Cruz; sc-74470), EME1 (Santa Cruz; sc-53275), KAP1 (Bethyl; A300-274A), P-KAP1 Ser 824 (Bethyl; A300-767A), β-tubulin (Cell Signaling; 86298S), histone H3 (Abcam; ab1791), CENPF (BD; 610768), Alexa-Fluor anti-rabbit 488 (ThermoFisher; A11070), Alexa-Fluor anti-mouse 594 (ThermoFisher; A11032), Alexa-Fluor anti-rabbit 594 (ThermoFisher; A-21207), Anti-rabbit HRP (Agilent; P0399), Anti-mouse HRP (Agilent; P0447).

### 2.4 Clonogenic survival assays

Appropriate cell lines were transfected with indicated siRNAs and/or treated with 1 µm AZD0156 (SelleckChem) for 24 hours where indicated. Seventy-two hours post transfection/treatment, cells were seeded at low density and treated with the denoted concentrations of Olaparib (AZD2281; Selleckchem) for 3 days. Olaparib was then replaced every 3 days for a total of 10 days. Colonies were then stained with 0.5 % crystal-violet (Sigma-Aldrich) in ddH^2^O and counted, or colonies were resolubilised using 1% SDS solution and absorbance measured at 595 nm. Data are expressed as percentage survival normalised to untreated controls for each condition.

### 2.5 Immunofluorescence

Permeabilisation, fixation and staining of cells on glass coverslips was carried out as previously described^22^. Images were acquired with a Nikon E600 Eclipse equipped with a 60x oil lens and Nikon Elements software, and foci numbers analysed using Image J (NIH). For each independent experiment > 100 nuclei were counted per condition.

### 2.6 Homologous recombination assay

Homologous recombination was measured using a CRISPR-based LMNA assay as previously described^39^. Briefly, HeLa cells were transfected with siRNA, seeded onto glass coverslips and co-transfected with plasmids encoding Cas9, a gRNA targeting LMNA, a donor plasmid encoding clover-LMNA, and a plasmid expressing RFP. Forty-eight hours later, coverslips were fixed using 4 % paraformaldehyde and cells permeabilised in PBS supplemented with Trition-X-100 (0.5 % v/v) for 5 minutes at room temperature. Coverslips were then blocked as above before being mounted onto slides and imaged as described above. Homologous recombination activity was quantified by enumerating clover-LMNA positive cells and normalising to RFP transfection efficiency. For each independent experiment > 100 nuclei were counted per condition.

### 2.7 Metaphase spreads

Cells were transfected with siRNA then treated with 10 µM Olaparib for 24 hours. Three hours prior to harvesting cells were treated with 0.1 µg/ml Colcemid (ThermoFisher Scientific). Cells were harvested by trypsinisation, incubated in prewarmed 0.075M KCl for 10 minutes, and fixed in ice cold fixative solution (3:1 methanol/acetic acid). Metaphases were dropped onto glass slides pre-soaked in fixative and placed on a heat block at 80 ° C for 1 minute, before being allowed to dry overnight. Slides were mounted with Duolink in situ mounting medium containing DAPI (Sigma-Aldrich) and imaged as above using a 100x oil lens. Radial chromosomes were enumerated using ImageJ software, and at least 50 metaphases were counted per condition for each independent experiment.

### 2.8 Chromatin fractionation

HeLa Kyoto iCas9 Cells were harvested by trypsination and lysed in chromatin fractionation buffer (20 mM Tris-HCl, 100 mM NaCl, 5 mM MgCl^2^, 10 % glycerol, 0.2 % IGEPAL CA-620, 0.5 mM DTT, protease inhibitor cocktail) on ice for 15 minutes. Lysates were clarified at 200 x g for 5 minutes at 4 °C, supernatant was removed to form a ‘soluble fraction’ and the chromatin pellet was washed and resuspended in fractionation buffer before two cycles of sonication at 10% power for 10 seconds.

### 2.9 Western blotting

Immunoblotting of whole cells extracts (WCE) or cellular fractions was performed as previously described^22,37^.

### 2.10 RNA sequencing

HeLa Kyoto iCas9 expressing SETD1A gRNA were transfected with control, ATM or BRCA1 siRNA as described above and Cas9 expression induced with doxycycline where appropriate. Total RNA was extracted from cell pellets using the RNeasy kit (Qiagen) with on-column DNase digest, and RNA quality confirmed using TapeStation (Aligent Technologies). mRNA was isolated using NEBNext Poly(A) mRNA magnetic isolation module and sequencing libraries were created using NEBNext Ultra II direction RNA library prep kit for Illumina (New England Biolabs). DNA libraries were validated with TapeStation (Aligent Technologies) and quantified using Qubit 2.0 Fluorometer (ThermoFisher Scientific), before being pooled and sequenced on Illumina NovaSeq 6000. Raw sequencing data was uploaded to the Galaxy web platform and analysed using the public server at https://usegalaxy.eu/^40^.

### 2.11 Quantitative polymerase chain reaction (qPCR)

cDNA was synthesised from 2 µg RNA (see above) using a High-Capacity cDNA Reverse Transcriptase kit (Applied Biosciences). cDNA was then quantified by TaqMan gene expression array (Applied Biosystems) with a Quantstudio 5 PCR system, and mRNA expression normalised to the levels of GUS1B.

### 2.12 Cancer genomics analysis

cBioPortal (https://www.cbioportal.org/)^41,42^ was used to access genomic data from TCGA lung adenocarcinoma, ovarian serous cystadenocarcinoma and the Metabric breast carcinoma datasets. Log2 mRNA expression of indicated genes was obtained from RNA-seq datasets. Patients were stratified based on SETD1A mRNA expression: Low= homozygous SETD1A deletion or mRNA expression <=-2 SD below mean; High= SETD1A mRNA expression >2 SD above mean. For Kaplan-Meir survival analysis *ATM*-, *BRCA1*- or *BRCA2*-deficient cancer patients were selected (defined as homozygous deletion, mutation, or mRNA expression <=-2 SD below mean) and overall survival data downloaded. Patients were then stratified by low (bottom 50% mRNA expression) or high SETD1A (upper 50% mRNA expression) gene expression.

### 2.13 ChIP-Seq analysis

Setd1a ChIP-Seq data from mouse embryonic stem cells was downloaded (GEO: GSE93538)^43^. Reads were aligned to mouse mm10 genome using Bowtie2, and peaks called using EaSeq. Plots were created in IGV Genome Browser for the selected genes.

### 2.14 Statistical analyses

Statistical significance of data was assessed using GraphPad Prism 10. Means were compared as indicated using either two-tailed unpaired Students t-test, one-way ANOVA or two-way ANOVA with post-hoc Tukeys correction for multiple comparisons. Kaplan-Meir survivals were analysed using a Log-rank (Mantel-Cox) test. Data points represent at least three independent biological repeats unless stated. In all cases * = *P* ≤ 0.05, ** = *P* ≤ 0.01, *** = *P* ≤ 0.001, **** = *P* ≤ 0.0001.

## 3. RESULTS

### 3.1 Loss of SETD1A reduces the sensitivity of ATM-deficient cells to Olaparib

Our previous work demonstrated that loss of the lysine methyltransferase SETD1A decreased the sensitivity of *BRCA1-*deficient cells to Olaparib or Talazoparib^22^, which we attributed to partial restoration of HR in the absence of SETD1A/H3K4me3, suggesting a new epigenetic mechanism of PARPi resistance^23^.

Multiple additional contexts of HR deficiency outside BRCA1 drive synthetic lethality with PARP inhibition, including alterations in *BRCA2, RAD51* and *ATM*^24,25,27,44^. However, the resistance mechanisms relevant to these other HR-deficient contexts are not well understood. To gain further insights into these mechanisms, we first assessed whether loss of SETD1A conveyed resistance to PARP inhibitors in the absence of ATM activity. To this end, we assessed the impact of Olaparib treatment on cells depleted of ATM and/or SETD1A (**Fig 1A-C** and **Fig S1A-B**). As expected, knockdown of ATM resulted in hypersensitivity to Olaparib treatment, in keeping with its role in promoting HR, whilst loss of SETD1A alone had no effect. Interestingly, the sensitivity of ATM-deficient cells to Olaparib was partially lost upon co-depletion of SETD1A. These findings were further validated in cells treated with an ATM inhibitor, AZD0156 (**Figs 1D-F**), which dramatically sensitised cells to Olaparib.

**Figure 1.**
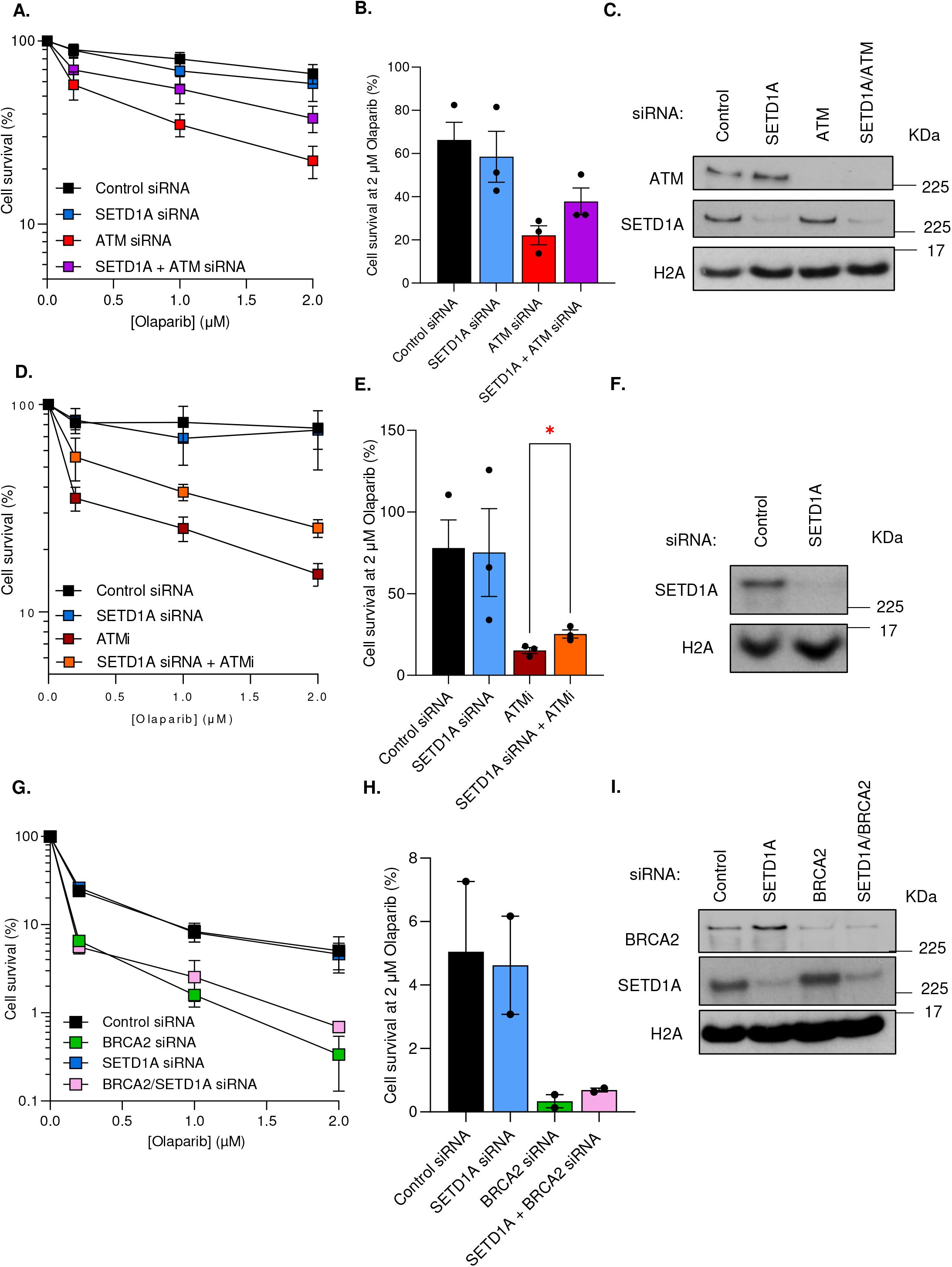
Loss of SETD1A reduces sensitivity of ATM-deficient cells to Olaparib. **(A-B)** HeLa cells were transfected with the indicated siRNAs and left to form colonies under continual exposure to Olaparib at the denoted doses for 10 days. Panel B denotes relative cell survival at 2 µM Olaparib. **(C)** Whole cell extracts of cells from A were analysed by immunoblotting using the indicated antibodies. **(D-E)** HeLa cells were transfected with the indicated siRNA, left for 48 hours, and treated with 1 µM of ATM inhibitor AZD0156. Cells were then left to form colonies for 10 days under continual exposure to Olaparib at the denoted doses. Panel E denotes relative cell survival at 2 µM Olaparib. **(F)** Whole cell extracts of cells from D were analysed by immunoblotting using the indicated antibodies. **(G-H)** MCF7 cells were transfected with the indicated siRNAs and left to form colonies under continual exposure to Olaparib at the denoted doses for 10 days. Panel H denotes relative cell survival at 2 µM Olaparib. **(I)** Whole cell extracts of cells from G were analysed by immunoblotting using the indicated antibodies. Data points in all cases represent mean±SEM from three independent biological repeats. * = P <0.05 as determined by an unpaired two-tailed Student’s -test.

PARP inhibitors are also approved in the treatment of *BRCA2*-deficient tumours: however, mechanisms of PARPi resistance differ substantially compared to cells lacking BRCA1^8,45^. For example, loss of pro NHEJ factors such as 53BP1 cannot rescue HR activity and induce PARPi resistance in *BRCA2*-mutated cells^11^. In keeping, no restorative effect on HR was observed upon SETD1A depletion in cells lacking BRCA2, demonstrating that SETD1A-mediated PARPi resistance is not universal across HR-deficient cell lines (**Figs 1G-I**). Together, these findings demonstrate that, in addition to *BRCA1*-deficient cells, sensitivity to PARP inhibition caused by loss of ATM function or its kinase activity can also be alleviated by depletion of SETD1A.

### 3.2 SETD1A depletion partially restores HR in ATM-deficient cells

In *BRCA1*-deficient cells, loss of pro-NHEJ factors such as 53BP1, RIF1, REV7-Shieldin, and the lysine methyltransferase SETD1A and its cofactor BOD1L restore HR activity^13-18,22,46^. We therefore hypothesised that the reduced PARPi sensitivity observed in ATM-deficient cells upon loss of SETD1A may also be due to restoration of HR.

To examine this, we first assessed RAD51 foci formation as a marker of HR following Olaparib treatment. Knockdown or inhibition of ATM resulted in a significant defect in RAD51 foci formation (**Figs 2A-C**). In both cases, depletion of SETD1A on these backgrounds increased RAD51 foci formation to near-normal levels, which was most notable in the presence of ATMi (**Fig 2C**). In agreement with our findings above, loss of SETD1A from BRCA2-depleted MCF-7 cells had no restorative effect on RAD51 foci formation (**Figs 2D-E**). To assess HR activity directly, we utilised a CRISPR-based assay in which cells utilise Clover-tagged Lamin as a template for functional HR, resulting in green, fluorescent ringed nuclei^39^. Using this assay, ATM knockdown or inhibition significantly decreased HR, whilst co-depletion of SETD1A from these cells resulted in partial HR restoration (**Fig 2E** and **Fig S1C**).

**Figure 2.**
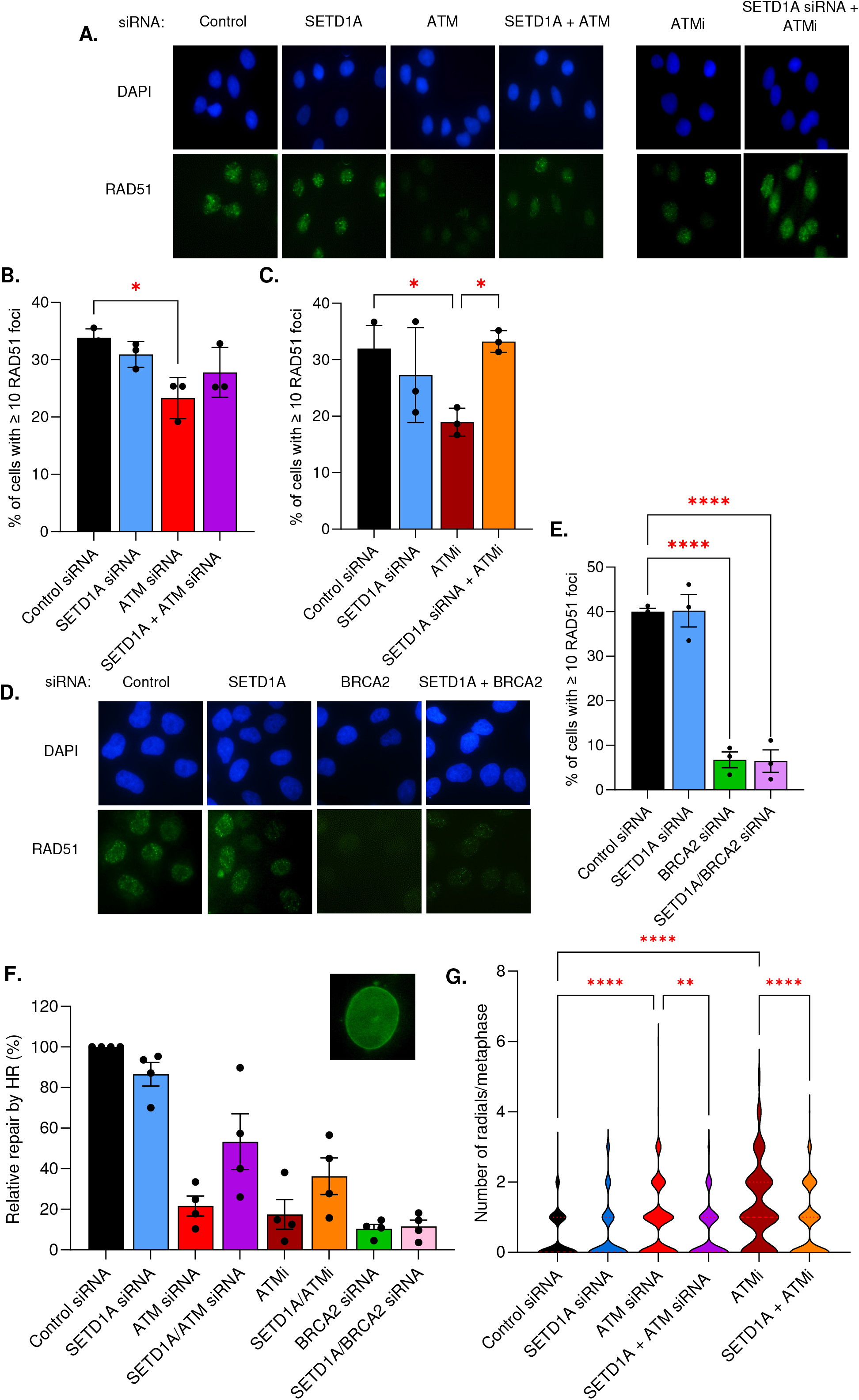
Loss of SETD1A partially restores homologous recombination (HR) activity in ATM-deficient cells treated with Olaparib. **(A-C)** HeLa cells were transfected with the indicated siRNAs and/or treated with 1 µM AZD0156 for 24 hours as indicated, and then treated with 5 µM Olaparib for a further 24 hours. Cells were immunostained with an antibody against RAD51 and nuclei counterstained with DAPI, and foci enumerated. Representative images are shown in A, and data from 3 independent biological repeats shown in B-C. **(D-E)** MCF-7 cells were transfected with the indicated siRNAs, and treated with 5 µM Olaparib for 24 hours. Cells were immunostained as in A: representative images are shown in D, and data from 3 independent biological repeats shown in E. **(F)** HeLa cells were transfected/treated as denoted, left for 24 h, and then transfected with CRISPR based LMNA-HR plasmids encoding Cas9 nickase, a fluorescent Lamin A/C and a plasmid encoding red fluorescent protein (RFP). Percentage HR was calculated by the number of fluorescent Lamin-expressing cells normalised to RFP expression. **(G)** Cells from F were incubated with 5 µM Olaparib for 24 hours, and the number of radial chromosomes per metaphase analysed by DAPI staining and quantified. In all cases data represent the mean ± SEM from at least three independent experiments. * = P < 0.05, ** = P < 0.01 ***= P < 0.0005, **** = P < 0.0001 as determined using a one-way ANOVA with post-hoc Tukey test for multiple comparisons.

PARP inhibition in HR-deficient cells is also characterised by the NHEJ-mediated formation of chromosomal radials which drive cell death. We therefore set out to confirm whether toxic end-joining was decreased following SETD1A depletion in ATM-deficient cells, using metaphase spreads to quantify radial chromosome formation. Consistent with the data above, cells lacking ATM displayed increased toxic radial formation upon Olaparib treatment, whilst co-depletion of SETD1A reduced this back to control levels (**Fig 2F** and **Fig S1D**). This provides further evidence that loss of SETD1A results in a shift in pathway usage from pro-NHEJ towards HR, thus restoring recombination in certain HR-deficient backgrounds.

In addition, the ability of PARPi to trap PARP1/2 onto damaged chromatin correlates with their potency *in vitro*^47^. We postulated that resistance to Olaparib upon loss of SETD1A might additionally reduce PARP1/2 trapping, leading to decreased cell death. Surprisingly however, levels of PARP1 on chromatin bore little correlation with sensitivity to PARPi (**Fig S2**): although both ATM deficiency and SETD1A deletion via Cas9 editing decreased PARP1 trapping after cell fractionation, they had opposite phenotypic effects. This strongly suggested that reduced PARP trapping was not responsible for PARPi sensitivity/resistance in these cells. Instead, our findings demonstrate that PARPi resistance upon loss of SETD1A arises due to loss of pro-NHEJ functions and restoration of HR.

### 3.3 Perturbing H3K4 methylation reduces sensitivity of ATM-deficient cells to Olaparib by partially restoring HR

SETD1A catalyses mono-, di- and tri-methylation of lysine 4 of histone H3 (H3K4), protecting stalled replication forks and promoting NHEJ^22,37,48^. To investigate whether H3K4me3 was directly responsible for our observations in *ATM*-deficient backgrounds, we used a mutant cell line expressing either wild-type GFP-tagged histone H3 or a H3K4A mutant which perturbs methylation at lysine 4, and that phenocopies loss of SETD1A^37,38^. Using these cells, we examined the impact of Olaparib treatment in the presence/absence of ATM. In agreement with our previous observations, ATM deficiency sensitised H3-WT-GFP cells to Olaparib and substantially compromised HR. In contrast, depletion of ATM only partially sensitised H3-K4A-GFP cells to PARPi (**Fig 3A-B**), and these cells showed increased levels of RAD51 foci (**Fig 3C-D**) and a reduction in the formation of toxic chromosomal radials (**Fig 3E**). In combination with previous findings, this suggests that loss of SETD1A-mediated H3K4 methylation influences PARPi sensitivity by restoring HR.

**Figure 3.**
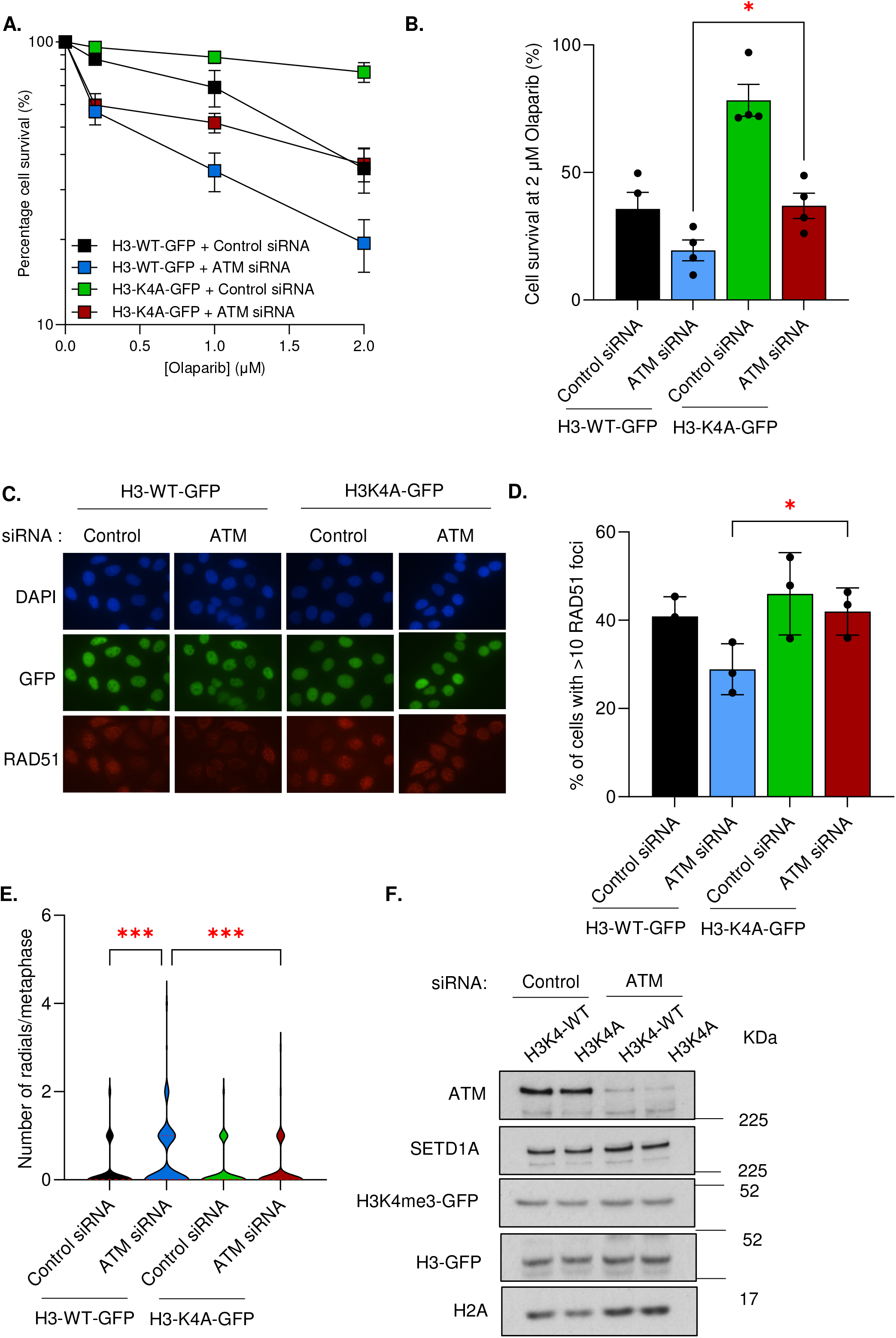
Loss of H3 lysine 4 methylation reduces sensitivity of ATM-deficient cells to Olaparib and partially restores homologous recombination. **(A-B)** H3K4-WT-GFP or H3K4A-GFP Hela cells were transfected with the indicated siRNAs and left to form colonies under continual exposure to Olaparib at the denoted doses for 10 days. Panel B denotes relative cell survival at 2 µM Olaparib. Data represent mean ± SEM from four independent repeats. **(C-D)** H3K4-WT-GFP or H3K4A-GFP cells from A were treated with 5 µM Olaparib for 24 hours, immunostained with antibodies against RAD51, and foci enumerated. Representative images are shown in C, and data from 3 independent biological repeats shown in D. **(E)** Cells from A were incubated with 5 µM Olaparib for 24 hours, and the number of radial chromosomes per metaphase analysed by DAPI staining and quantified. **(F)** Whole cell extracts of cells from A were analysed by immunoblotting using the indicated antibodies. In all cases, data represent mean ± SEM from three biologically independent experiments. * = P < 0.05 as determined by n unpaired two-tailed Students t-test (B, D), ***= P < 0.0005 as determined by a one-way ANOVA with post-hoc Tukey test for multiple comparisons (E).

### 3.4 Loss of SETD1A reduces sensitivity of HR-deficient cancer cell lines to Olaparib

We next wanted to extend our findings from HeLa cells into cancer cell lines bearing pathogenic *BRCA1* or *ATM* mutations. We therefore selected 3 cell lines encoding *BRCA1* (UWB1.89 and HCC1937) or *ATM* (H1395) mutations, with non-mutated SKOV3, MCF-7 and A549 cell lines as cancer-type matched controls. As expected, *BRCA1*-mutated UWB1.289 ovarian cancer (**Fig 4A-B**) and HCC1937 breast cancer cells (**Fig 4C-D**) displayed significant sensitivity to Olaparib, whilst *BRCA1*-proficient SKOV3 ovarian or MCF-7 breast cancer cells did not. In both cases, depletion of SETD1A reduced the sensitivity of *BRCA1*-mutated cell lines to Olaparib but had no effect on *BRCA1* wild-type cells. Similar results were obtained in *ATM*-mutated H1395 non-small cell lung cells, where SETD1A depletion again compromised PARPi sensitivity (**Fig 4E-F**). Moreover, and in agreement with preceding data, loss of SETD1A also restored Olaparib-induced RAD51 foci formation and thus HR activity in both *BRCA1*-mutated ovarian and breast cancer cell lines and in *ATM*-mutated lung cancer cells (**Fig 4G-I**). In concert, these findings firmly establish that loss of SETD1A drives PARPi resistance in breast, ovarian and lung cancer cells by restoring HR activity.

**Figure 4.**
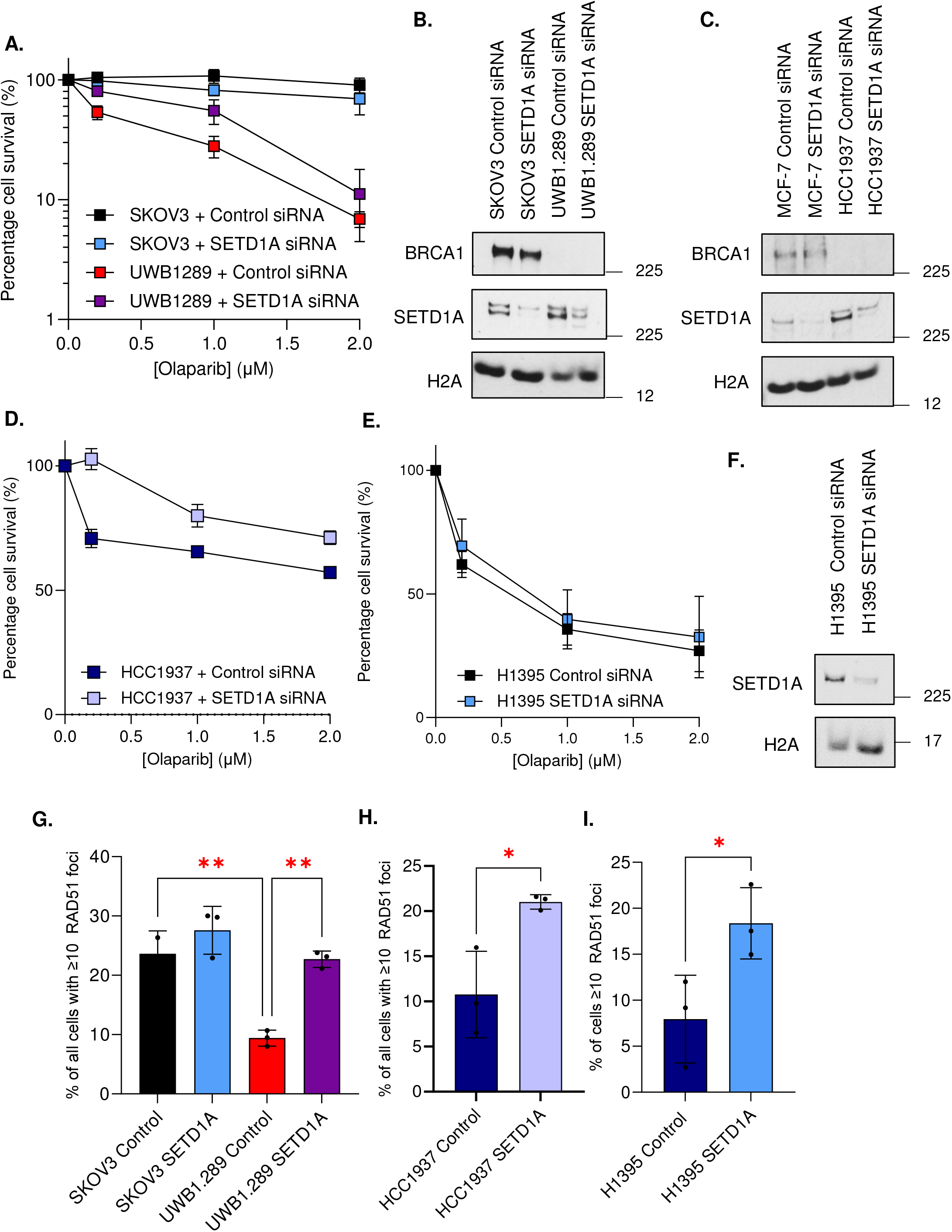
Loss of SETD1A reduces sensitivity of HR-deficient cancer cell lines to Olaparib via partial restoration of HR. **(A-F)** SKOV3, UWB1.289, MCF-7, HCC1937 or H1395 cells were transfected with the indicated siRNAs. Cells were plated at low density and left to form colonies under continual exposure to Olaparib at the denoted doses for 10 days (panels A, D, E). Whole cell extracts were analysed by immunoblotting in panels B, C and F. **(G-I)** Transfected cells from A, D and E were treated with 5 µM Olaparib for 24 hours, and immunostained with antibodies against RAD51 and CENPF. RAD51 foci in G1 (CENPF negative) or G2 (CENPF positive) cells were enumerated and shown in G-I. All data represent the mean ± SEM from >3 independent experiments. * = P < 0.05, ** = P < 0.01 ***= P < 0.0005, **** = P < 0.0001 as determined a one-way ANOVA with post-hoc Tukey test for multiple comparisons.

### 3.5 SETD1A-dependent transcription of EME1 drives PARPi resistance

Given that SETD1A-mediated H3K4me plays a key role in transcriptional regulation^48^, we postulated that gene expression changes may account for our observations. To investigate this, we established an isogenic HeLa cell system that expressed a gRNA targeting the *SETD1A* locus, in which SETD1A could be deleted by inducing Cas9 expression (HeLa Kyoto iCas9; **Fig S3**). We then depleted either ATM or BRCA1 and performed RNA sequencing (RNA-seq) analysis to determine differentially expressed (DEGs) regulated by SETD1A (**Fig 5A-B**). Surprisingly, loss of SETD1A had a limited impact on gene expression in HR-proficient cells, with only 13 transcripts showing significant deregulation upon Cas9 expression. Next, to identify genes deregulated in PARPi-sensitive or resistant contexts, we compared DEGs between *BRCA1*-deficient, *ATM*-deficient, and HR-proficient backgrounds in the presence/absence of SETD1A. This identified 5 genes whose expression was commonly regulated by SETD1A irrespective of HR proficiency (denoted in white in **Fig 5C**), and one gene, EME1, that was downregulated in different HR-deficient PARPi-resistance backgrounds (indicated in yellow in **Fig 5C**). Importantly, these gene expression changes were validated using qPCR in additional samples from each cell type, except for PEX12 (**Fig S4**). By analysing mouse ChIP-seq datasets^43^, we also observed that SETD1A was highly enriched at the TSS of EME1, but not at the TSS of genes regulated by BRCA1 or ATM (**Fig 5D** and **Fig S5**). Finally, we confirmed our findings at the protein level, where SETD1A deletion substantially decreased EME1 protein expression (**Fig 5E**). Overall, these analyses suggests that whilst the transcription of only a small number of genes is dependent on SETD1A in our iCas9 model, down-regulation of EME1 might represent a common mechanism linked to PARPi resistance.

**Figure 5.**
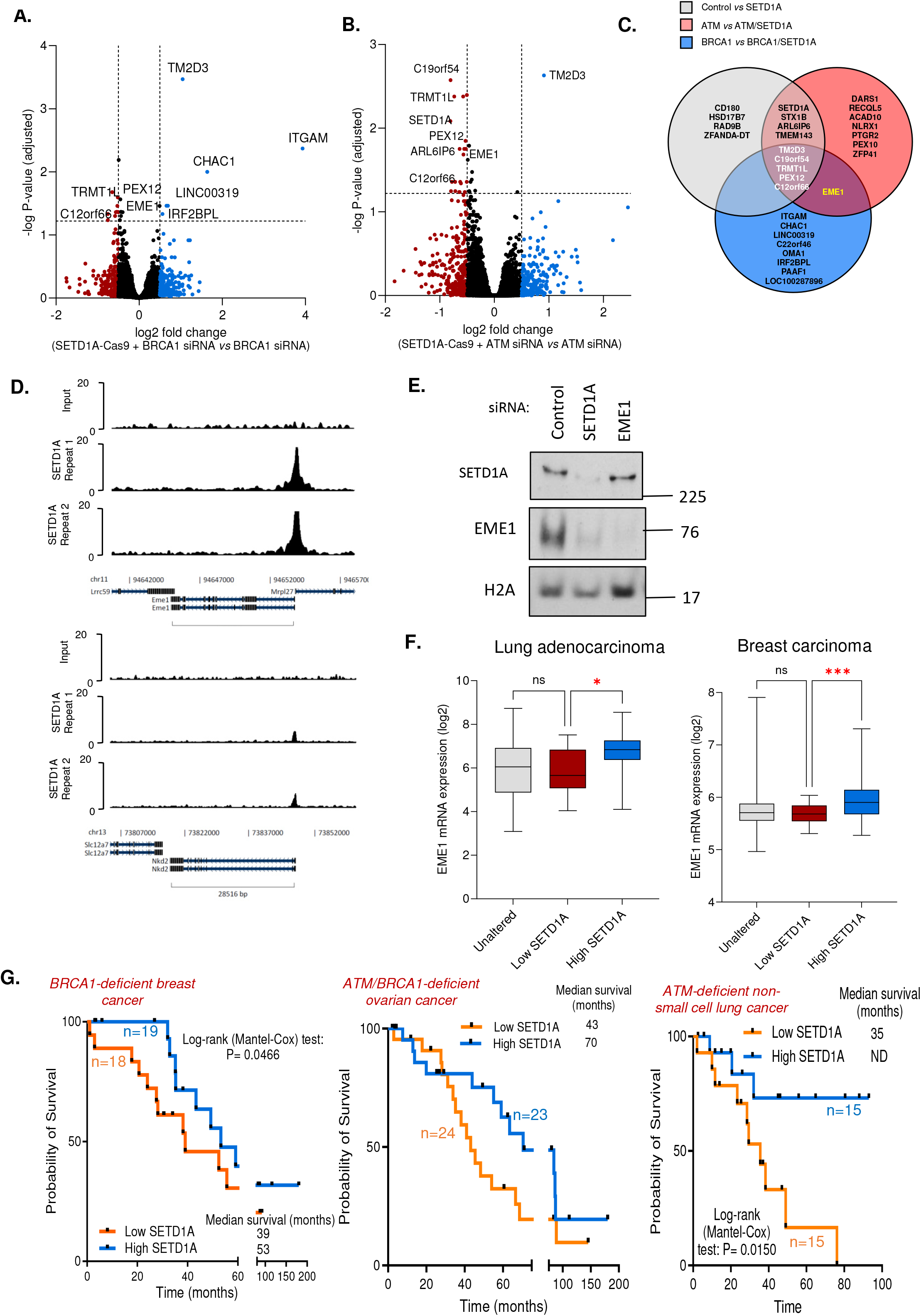
SETD1A-dependent EME1 transcription in ATM- and BRCA1-deficient cells. **(A-B)** Inducible Hela Kyoto iCas9 cells expressing SETD1A gRNA were treated with doxycycline and transfected with the indicated siRNAs. Gene expression was analysed using RNA-seq and differential gene expression displayed as a volcano plot. Significantly downregulated genes are represented in red and significantly upregulated genes denoted in blue. Cutoff values for differentially expressed genes were Log2 fold change = 0.5 and -log (adjusted P-value) = 1.22. Data points represent means from 3 independent repeats. **(C)** Venn diagram of overlapping differentially expressed genes from A-B. **(D)** Chromatin immunoprecipitation profiles of murine Setd1a over the TSS of genes from C. Source data are from ^43^. **(E)** Whole cell extracts of Hela Kyoto iCas9 cells expressing SETD1A gRNA were analysed by immunoblotting using the denoted antibodies. **(F)** Log2 mRNA expression of EME1 in lung adenocarcinoma or breast carcinoma patients triaged by SETD1A expression. Low= homozygous SETD1A deletion or mRNA expression <=-2 SD below mean; high= SETD1A mRNA expression >2 SD above mean. **(G)** Kaplan-Meir curves denoting overall survival for the indicated cancer patient subtypes stratified by low (lower 50%) or high (upper 50%) SETD1A mRNA expression. ****= P < 0.0001 as determined by a two-tailed unpaired Students t-test (panel E). * = P <0.05, ***= P < 0.0005, as determined by a one-way ANOVA with post-hoc Tukey test for multiple comparisons (panel F).Statistical significance of the data in G was determined by a Log-rank (Mantel-Cox) test.

### 3.6 Deregulated gene expression in cancer patients with altered SETD1A expression

We next set out to investigate the relevance of our findings using cancer genomics data^41,42^. We first stratified lung adenocarcinoma, breast carcinoma or ovarian carcinoma patients by SETD1A mRNA expression and then assessed levels of each DEG. Consistent with our previous analyses, patients with low SETD1A levels (<-2 SD log2 mRNA expression) had significantly decreased expression of genes identified as downregulated by RNA-seq, including EME1 (**Fig 5F** and **Fig S6**). We next investigated whether these observations affected clinical outcome by examining overall survival. Firstly, we identified cohorts of HR-deficient breast, ovarian and lung cancer patients^41,42^, stratified them by high/low SETD1A mRNA expression as before, and analysed overall survival rates. This revealed that *BRCA1*- and *ATM*-deficient cancer patients with low SETD1A mRNA expression had poorer overall survival outcomes compared to those with higher levels of SETD1A (**Fig 5G-I**). This was not the case for BRCA2-deficient patients, nor was it the case for HR-proficient patients (**Fig S7A**). Finally, we assessed whether EME1 might correlate with patient survival in a similar way to SETD1A. Interestingly, in most cases, EME1 expression did not correlate with increased/decreased patient survival, except in the case of BRCA1-deficient breast cancer, where low EME1 expression associated with a favourable prognosis (**Fig S7C-F**). Thus, SETD1A, and to a limited extent EME1, correlates with survival of HR-deficient patients.

### 3.7 Loss of EME1 drives PARPi resistance in HR-deficient cells by restoring HR

Given that SETD1A promotes EME1 transcription *in vitro*, and that this correlation was observed in cancer genomic datasets, we hypothesised that EME1 downregulation might drive PARPi resistance observed in the absence of SETD1A. To assess this, we depleted EME1 and/or BRCA1/ATM from HeLa cells (**Fig 6A**) and assessed the response to Olaparib. Excitingly, EME1 loss phenotypically mirrored SETD1A depletion in both these backgrounds *i*.*e*. its loss compromised PARPi sensitivity (**Fig 6B-E**). Surprisingly, given its known roles in later stages of HR where it processes recombination intermediates^49-51^, loss of EME1 also restored levels of HR in both ATM- or BRCA1-deficient cells after exposure to PARPi (**Fig 6F-G**), precisely as observed with SETD1A loss. Furthermore, downregulation of EME1 was also observed when SETD1A was depleted from BRCA1-mutated HCC1937 breast cancer cells, or ATM-mutated H1395 lung cancer cells (**Fig 6H**), suggesting that this represents a conserved mechanism in multiple cancer types. Together, this demonstrates that loss of SETD1A-dependent EME1 transcription drives PARPi resistance in HR-deficient cells.

**Figure 6.**
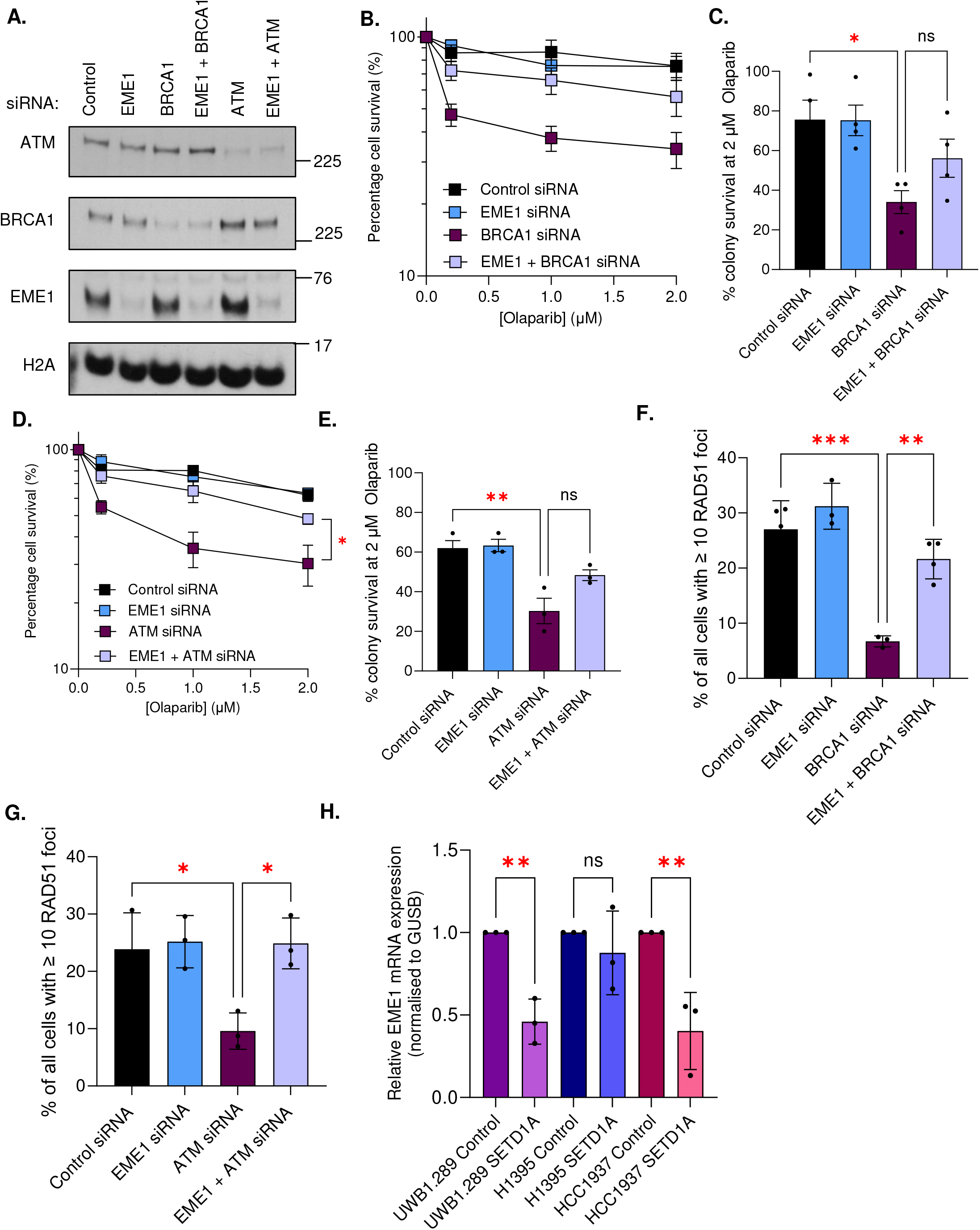
Deregulated EME1 expression partially restores HR and drives resistance of Olaparib in ATM- and BRCA1-deficient cells. **(A-E)** HeLa cells were transfected with the indicated siRNAs for 48h. Whole cell extracts were analysed by immunoblotting (A). Cells were plated at low density and left to form colonies under continual exposure to Olaparib at the denoted doses for 10 days (B and D). Panels C and E denotes relative cell survival at 2 µM Olaparib. **(F-G)** Cells from A were treated with 5 µM Olaparib for 24 hours, immunostained with antibodies against RAD51, and foci enumerated. **(H)** mRNA was isolated form cells from A, and EME1 and GUSB mRNA expression levels determined by qPCR. In all cases, data points represent the mean ± SEM from 3 independent repeats. * = P <0.05, ** = P <0.01, ***= P < 0.0005, ****= P < 0.0001 as determined by a two-tailed unpaired Students t-test.

In summary, we show that SETD1A loss renders *ATM*-deficient and *BRCA1*-mutated breast, lung, and ovarian cancer cells resistant to PARPi by restoring HR, and that this is partly driven by defective EME1 transcription.

## 4. DISCUSSION

Multiple mechanisms of PARPi resistance in BRCA-deficient cells have now been described and have highlighted clear differences between resistance arising in *BRCA1*- and *BRCA2*-mutated cancers ^e.g.8,52,53^. However, to date, there have been few studies examining PARPi resistance in cells lacking other HR factors. This, coupled with mixed reports of PARPi efficacy in trials in other cancer backgrounds of HR-deficiency^31,33-36,54-57^, led us to examine whether loss of the lysine methyltransferase and pro-NHEJ factor SETD1A influenced PARPi efficacy outside BRCA1.

Here we show that loss of SETD1A also confers PARPi resistance in *ATM*-deficient cancer cells. Inhibition of ATM activity or depletion of ATM sensitised cervical cancer lines to Olaparib, in line with numerous previous studies demonstrating that ATM deficiency correlates with PARPi sensitivity^24-27,58-61^. However, depletion of SETD1A rendered these cells at least partially resistant to PARPi (**Fig 1**). Moreover, we observed similar resistance in *ATM*-mutated H1395 NSCLC cells depleted of SETD1A (**Fig 4**), demonstrating that this phenomenon was not cell-type specific or associated with ATM inhibitor or depletion using siRNA. Although the extent of PARP1 trapping by Olaparib was not affected by SETD1A depletion in these models (**Fig S2**), we observed restoration of HR by RAD51 foci, Cas9-nickase mediated recombination and decreased toxic chromosome radials (**Figs 2** and **4**). This is entirely consistent with our previous findings that SETD1A depletion restored HR in *BRCA1*-deficient cells^22,23^. It also suggests that a deficiency in resection caused by loss of ATM can be overcome by compromising the 53BP1-RIF1-Shieldin pathway. This agrees with studies in *ATM-*deficient breast cancer cells showing that 53BP1 depletion causes PARPi resistance^46^, but is at odds with the demonstration that loss of 53BP1 cannot suppress the PARPi sensitivity of Atm^*-/-*^ mESCs^62^. Whilst it is difficult to reconcile these differences, it may be that the genetic background and/or tumoral nature of MCF-7, CAL51, HeLa or H1395 cells used in our study or that of Hong and colleagues^46^ is responsible. In addition, the necessity for ATM in activating the 53BP1-RIF1 axis *via* 53BP1 phosphorylation likely plays an important role^13,15^. Nevertheless, our findings strongly suggest that perturbation of pro-NHEJ pathways by loss of SETD1A drives PARPi resistance.

Our findings also demonstrate that loss of SETD1A represents a conserved mechanism of PARPi resistance across *BRCA1-* and *ATM-*deficient cancer cells (this study and^22^) but this is not extended to cells lacking *BRCA2* (**Figs 1** and **2**). Moreover, the survival of *BRCA1-* or *ATM-*deficient cancer patients could be triaged by SETD1A expression, which was not the case for those lacking functional *BRCA2*, nor those with functional copies/levels of these genes (**Fig S6**). These observations are likely due to the differing roles of ATM, BRCA1 and BRCA2 in HR. Whilst ATM and BRCA1 promote the initial steps of HR and resection and their loss can partially be overcome by loss of competing pathways, BRCA2 is absolutely required for RAD51-dependent recombination. This is in keeping with multiple studies demonstrating that loss of pro-NHEJ factors cannot drive PARPi resistance in *BRCA2*-deficient cells^52,53^.

Interestingly, SETD1A-dependent methylation of H3K4 and subsequent transcription also seems to underpin PARPi resistance in HR-deficient cells lacking this KMT. Firstly, perturbation of H3K4me using our H3-K4A-GFP system phenocopied SETD1A loss, alleviating PARPi sensitivity, restoring HR and decreasing PARPi-induced toxic radial formation in the absence of ATM (**Fig 3**). Moreover, RNAseq analyses revealed that expression of EME1 mRNA was tightly correlated with PARPi-resistance, with SETD1A-depleted cells displaying ∼2-fold reduction in EME1 mRNA and protein levels (**Figs 5** and **S5**). In addition, SETD1A was enriched at the TSS of EME1 in mESCs but was absent from genes regulated by either BRCA1 or ATM, suggesting both specificity and a wider role for SETD1A in transcriptionally regulating EME1. Moreover, loss/perturbation of EME1 phenocopies SETD1A loss in the absence of both BRCA1 and ATM and led to the restoration of HR s (**Fig 6**). EME1 forms part of the SLX-MUS81-EME1 structure-specific nuclease that plays a key role in resolving Holliday junctions, including during the latter stages of HR^43,49-51,63^. Our findings are thus in broad agreement with findings from the D’Andrea lab^64^ that loss of MUS81 renders BRCA2-deficient cells resistant to PARPi. However, our data also suggest a new anti-recombinogenic role for EME1 in HR-deficient cells. Whether this involves other members of this nuclease complex, and whether the replication stress-specific paralog EME2^65^ is also involved, remains to be determined.

Previously, we attributed the PARPi resistance in *BRCA1*-deficient cells lacking SETD1A to a failure to recruit RIF1, arising from defective H3K4 methylation and a subsequent loss of binding between H3K4me3 and RIF1 at damaged sites^22^. Here, we offer an additional mechanism controlled by SETD1A/H3K4me that influences PARPi sensitivity: the transcription of EME1. Although this appears contradictory, this mirrors the multiple functions of other DNA repair proteins that influence PARPi response, for example the roles of BRCA1 in HR, fork protection, and ssDNA gap formation^8-10,66,67^. The respective contributions of these mechanisms to PARPi resistance, or whether these SETD1A-dependent mechanisms are interlinked, are clearly areas for future study.

Finally, these findings reinforce our previous speculation^23^ that SETD1A status/activity could be a useful prognostic biomarker to monitor/predict PARPi response. Indeed, SETD1A expression correlates with overall survival in HR-deficient breast, ovarian and lung cancers (**Fig 5**). It is plausible that monitoring SETD1A levels or using EME1 expression or H3K4me3 as proxies for its activity might be useful to predict PARPi resistance, although this warrants closer investigation. It is tempting to speculate that this may ultimately improve the use of PARPi such as Olaparib in cancers where its efficacy has been limited (e.g. NSCLC^36^), by identifying patients amenable for therapy.

## ACKNOWLEDGEMENTS

We thank Clare Davies, Tatjana Stankovic, Marco Saponaro and other members of the Birmingham Centre for Genome Biology (BCGB) for invaluable discussions. We are grateful to Simon Boulton for providing iCas9 cells and lentiviruses, and to Graham Dellaire for providing plasmids for CRISPR-based HR assays^39^.

## AUTHORS’ CONTRIBUTIONS

Experiments were designed by E.S., R. B., and M.R.H. Experiments and data analysis were carried out by E.S., R. B., and R.S. M.R.H. and E.S. wrote the paper with contributions and editing by all other authors.

## FUNDING INFORMATION

E.S. was funded by the University of Birmingham and Cancer Research UK (C17422/A25154). R.B. was funded by a Birmingham Fellowship (204846/Z/16/Z) awarded to M.R.H. by the University of Birmingham. M.R.H. was funded by a Birmingham Fellowship (204846/Z/16/Z), a Medical Research Council Career Development Fellowship (MR/P009085/1), and a UKRI MRC Impact Acceleration (MR/X502996/1).

## Notes

**Conflict of interest**: The authors declare no conflict of interest.

### Competing Interest Statement

The authors have declared no competing interest.

## REFERENCES

1. Farmer H, McCabe N, Lord CJ, Tutt AN, Johnson DA, Richardson TB et al. Targeting the DNA repair defect in BRCA mutant cells as a therapeutic strategy. Nature 2005; 434(7035): 917–921; doi 10.1038/nature03445.

2. Lord CJ, Ashworth A. PARP inhibitors: Synthetic lethality in the clinic. Science 2017; 355(6330): 1152–1158; e-pub ahead of print 2017/03/18; doi 10.1126/science.aam7344.

3. Robson M, Im SA, Senkus E, Xu B, Domchek SM, Masuda N et al. Olaparib for Metastatic Breast Cancer in Patients with a Germline BRCA Mutation. N Engl J Med 2017; 377(6): 523-533; e-pub ahead of print 20170604; doi 10.1056/NEJMoa1706450.

4. Moore K, Colombo N, Scambia G, Kim BG, Oaknin A, Friedlander M et al. Maintenance Olaparib in Patients with Newly Diagnosed Advanced Ovarian Cancer. N Engl J Med 2018; 379(26): 2495–2505; e-pub ahead of print 20181021; doi 10.1056/NEJMoa1810858.

5. Golan T, Hammel P, Reni M, Van Cutsem E, Macarulla T, Hall MJ et al. Maintenance Olaparib for Germline BRCA-Mutated Metastatic Pancreatic Cancer. N Engl J Med 2019; 381(4): 317-327; e-pub ahead of print 20190602; doi 10.1056/NEJMoa1903387.

6. Abida W, Patnaik A, Campbell D, Shapiro J, Bryce AH, McDermott R et al. Rucaparib in Men With Metastatic Castration-Resistant Prostate Cancer Harboring a BRCA1 or BRCA2 Gene Alteration. J Clin Oncol 2020; 38(32): 3763–3772; e-pub ahead of print 20200814; doi 10.1200/JCO.20.01035.

7. de Bono J, Mateo J, Fizazi K, Saad F, Shore N, Sandhu S et al. Olaparib for Metastatic Castration-Resistant Prostate Cancer. N Engl J Med 2020; 382(22): 2091–2102; e-pub ahead of print 20200428; doi 10.1056/NEJMoa1911440.

8. Noordermeer SM, van Attikum H. PARP Inhibitor Resistance: A Tug-of-War in BRCA-Mutated Cells. Trends Cell Biol 2019; 29(10): 820–834; e-pub ahead of print 20190814; doi 10.1016/j.tcb.2019.07.008.

9. Dias MP, Moser SC, Ganesan S, Jonkers J. Understanding and overcoming resistance to PARP inhibitors in cancer therapy. Nat Rev Clin Oncol 2021; 18(12): 773–791; e-pub ahead of print 20210720; doi 10.1038/s41571-021-00532-x.

10. Kyo S, Kanno K, Takakura M, Yamashita H, Ishikawa M, Ishibashi T et al. Clinical Landscape of PARP Inhibitors in Ovarian Cancer: Molecular Mechanisms and Clues to Overcome Resistance. Cancers (Basel) 2022; 14(10); e-pub ahead of print 20220519; doi 10.3390/cancers14102504.

11. Bouwman P, Aly A, Escandell JM, Pieterse M, Bartkova J, van der Gulden H et al. 53BP1 loss rescues BRCA1 deficiency and is associated with triple-negative and BRCA-mutated breast cancers. Nat Struct Mol Biol 2010; 17(6): 688–695; e-pub ahead of print 20100509; doi 10.1038/nsmb.1831.

12. Jaspers JE, Kersbergen A, Boon U, Sol W, van Deemter L, Zander SA et al. Loss of 53BP1 causes PARP inhibitor resistance in Brca1-mutated mouse mammary tumors. Cancer Discov 2013; 3(1): 68–81; e-pub ahead of print 20121025; doi 10.1158/2159-8290.CD-12-0049.

13. Chapman JR, Barral P, Vannier JB, Borel V, Steger M, Tomas-Loba A et al. RIF1 is essential for 53BP1-dependent nonhomologous end joining and suppression of DNA double-strand break resection. Mol Cell 2013; 49(5): 858–871; e-pub ahead of print 2013/01/22; doi 10.1016/j.molcel.2013.01.002.

14. Dev H, Chiang TW, Lescale C, de Krijger I, Martin AG, Pilger D et al. Shieldin complex promotes DNA end-joining and counters homologous recombination in BRCA1-null cells. Nat Cell Biol 2018; 20(8): 954–965; e-pub ahead of print 2018/07/20; doi 10.1038/s41556-018-0140-1.

15. Escribano-Diaz C, Orthwein A, Fradet-Turcotte A, Xing M, Young JT, Tkac J et al. A cell cycle-dependent regulatory circuit composed of 53BP1-RIF1 and BRCA1-CtIP controls DNA repair pathway choice. Mol Cell 2013; 49(5): 872–883; e-pub ahead of print 2013/01/22; doi 10.1016/j.molcel.2013.01.001.

16. Noordermeer SM, Adam S, Setiaputra D, Barazas M, Pettitt SJ, Ling AK et al. The shieldin complex mediates 53BP1-dependent DNA repair. Nature 2018; 560(7716): 117–121; doi 10.1038/s41586-018-0340-7.

17. Xu G, Chapman JR, Brandsma I, Yuan J, Mistrik M, Bouwman P et al. REV7 counteracts DNA double-strand break resection and affects PARP inhibition. Nature 2015; 521(7553): 541–544; e-pub ahead of print 20150323; doi 10.1038/nature14328.

18. Zimmermann M, Lottersberger F, Buonomo SB, Sfeir A, de Lange T. 53BP1 regulates DSB repair using Rif1 to control 5’ end resection. Science 2013; 339(6120): 700–704; e-pub ahead of print 2013/01/12; doi 10.1126/science.1231573.

19. Castroviejo-Bermejo M, Cruz C, Llop-Guevara A, Gutierrez-Enriquez S, Ducy M, Ibrahim YH et al. A RAD51 assay feasible in routine tumor samples calls PARP inhibitor response beyond BRCA mutation. EMBO Mol Med 2018; 10(12); e-pub ahead of print 2018/11/01; doi 10.15252/emmm.201809172.

20. Pellegrino B, Herencia-Ropero A, Llop-Guevara A, Pedretti F, Moles-Fernandez A, Viaplana C et al. Preclinical In Vivo Validation of the RAD51 Test for Identification of Homologous Recombination-Deficient Tumors and Patient Stratification. Cancer Res 2022; 82(8): 1646-1657; e-pub ahead of print 2022/04/16; doi 10.1158/0008-5472.CAN-21-2409.

21. Waks AG, Cohen O, Kochupurakkal B, Kim D, Dunn CE, Buendia Buendia J et al. Reversion and non-reversion mechanisms of resistance to PARP inhibitor or platinum chemotherapy in BRCA1/2-mutant metastatic breast cancer. Ann Oncol 2020; 31(5): 590–598; e-pub ahead of print 20200220; doi 10.1016/j.annonc.2020.02.008.

22. Bayley R, Borel V, Moss RJ, Sweatman E, Ruis P, Ormrod A et al. H3K4 methylation by SETD1A/BOD1L facilitates RIF1-dependent NHEJ. Mol Cell 2022; 82(10): 1924–1939 e1910; e-pub ahead of print 2022/04/20; doi 10.1016/j.molcel.2022.03.030.

23. Bayley R, Sweatman E, Higgs MR. New perspectives on epigenetic modifications and PARP inhibitor resistance in HR-deficient cancers. Cancer Drug Resist 2023; 6(1): 35–44; e-pub ahead of print 20230104; doi 10.20517/cdr.2022.73.

24. Abbotts R, Dellomo AJ, Rassool FV. Pharmacologic Induction of BRCAness in BRCA-Proficient Cancers: Expanding PARP Inhibitor Use. Cancers (Basel) 2022; 14(11); e-pub ahead of print 20220526; doi 10.3390/cancers14112640.

25. McCabe N, Turner NC, Lord CJ, Kluzek K, Bialkowska A, Swift S et al. Deficiency in the repair of DNA damage by homologous recombination and sensitivity to poly(ADP-ribose) polymerase inhibition. Cancer Res 2006; 66(16): 8109–8115; doi 10.1158/0008-5472.CAN-06-0140.

26. Murai J, Pommier Y. BRCAness, Homologous Recombination Deficiencies, and Synthetic Lethality. Cancer Res 2023; 83(8): 1173–1174; doi 10.1158/0008-5472.CAN-23-0628.

27. Parvin S, Akter J, Takenobu H, Katai Y, Satoh S, Okada R et al. ATM depletion induces proteasomal degradation of FANCD2 and sensitizes neuroblastoma cells to PARP inhibitors. BMC Cancer 2023; 23(1): 313; e-pub ahead of print 20230405; doi 10.1186/s12885-023-10772-y.

28. Choi M, Kipps T, Kurzrock R. ATM Mutations in Cancer: Therapeutic Implications. Mol Cancer Ther 2016; 15(8): 1781–1791; e-pub ahead of print 20160713; doi 10.1158/1535-7163.MCT-15-0945.

29. Renwick A, Thompson D, Seal S, Kelly P, Chagtai T, Ahmed M et al. ATM mutations that cause ataxia-telangiectasia are breast cancer susceptibility alleles. Nat Genet 2006; 38(8): 873–875; e-pub ahead of print 20060709; doi 10.1038/ng1837.

30. Bryant HE, Helleday T. Inhibition of poly (ADP-ribose) polymerase activates ATM which is required for subsequent homologous recombination repair. Nucleic Acids Res 2006; 34(6): 1685–1691; e-pub ahead of print 20060323; doi 10.1093/nar/gkl108.

31. Weston VJ, Oldreive CE, Skowronska A, Oscier DG, Pratt G, Dyer MJ et al. The PARP inhibitor olaparib induces significant killing of ATM-deficient lymphoid tumor cells in vitro and in vivo. Blood 2010; 116(22): 4578–4587; e-pub ahead of print 2010/08/27; doi 10.1182/blood-2010-01-265769.

32. Mateo J, Carreira S, Sandhu S, Miranda S, Mossop H, Perez-Lopez R et al. DNA-Repair Defects and Olaparib in Metastatic Prostate Cancer. N Engl J Med 2015; 373(18): 1697–1708; doi 10.1056/NEJMoa1506859.

33. Bang YJ, Im SA, Lee KW, Cho JY, Song EK, Lee KH et al. Randomized, Double-Blind Phase II Trial With Prospective Classification by ATM Protein Level to Evaluate the Efficacy and Tolerability of Olaparib Plus Paclitaxel in Patients With Recurrent or Metastatic Gastric Cancer. J Clin Oncol 2015; 33(33): 3858–3865; e-pub ahead of print 20150817; doi 10.1200/JCO.2014.60.0320.

34. Schmitt A, Knittel G, Welcker D, Yang TP, George J, Nowak M et al. ATM Deficiency Is Associated with Sensitivity to PARP1-and ATR Inhibitors in Lung Adenocarcinoma. Cancer Res 2017; 77(11): 3040–3056; e-pub ahead of print 2017/04/02; doi 10.1158/0008-5472.CAN-16-3398.

35. Karmokar A, Sargeant R, Hughes AM, Baakza H, Wilson Z, Talbot S et al. Relevance of ATM Status in Driving Sensitivity to DNA Damage Response Inhibitors in Patient-Derived Xenograft Models. Cancers (Basel) 2023; 15(16); e-pub ahead of print 20230821; doi 10.3390/cancers15164195.

36. Ramalingam SS, Novello S, Guclu SZ, Bentsion D, Zvirbule Z, Szilasi M et al. Veliparib in Combination With Platinum-Based Chemotherapy for First-Line Treatment of Advanced Squamous Cell Lung Cancer: A Randomized, Multicenter Phase III Study. J Clin Oncol 2021; 39(32): 3633–3644; e-pub ahead of print 20210826; doi 10.1200/JCO.20.03318.

37. Higgs MR, Sato K, Reynolds JJ, Begum S, Bayley R, Goula A et al. Histone Methylation by SETD1A Protects Nascent DNA through the Nucleosome Chaperone Activity of FANCD2. Mol Cell 2018; 71(1): 25–41 e26; e-pub ahead of print 2018/06/26; doi 10.1016/j.molcel.2018.05.018.

38. Sato K, Ishiai M, Toda K, Furukoshi S, Osakabe A, Tachiwana H et al. Histone chaperone activity of Fanconi anemia proteins, FANCD2 and FANCI, is required for DNA crosslink repair. EMBO J 2012; 31(17): 3524–3536; e-pub ahead of print 20120724; doi 10.1038/emboj.2012.197.

39. Pinder J, Salsman J, Dellaire G. Nuclear domain ‘knock-in’ screen for the evaluation and identification of small molecule enhancers of CRISPR-based genome editing. Nucleic Acids Res 2015; 43(19): 9379–9392; e-pub ahead of print 2015/10/03; doi 10.1093/nar/gkv993.

40. Galaxy C. The Galaxy platform for accessible, reproducible and collaborative biomedical analyses: 2022 update. Nucleic Acids Res 2022; 50(W1): W345-W351; doi 10.1093/nar/gkac247.

41. Cerami E, Gao J, Dogrusoz U, Gross BE, Sumer SO, Aksoy BA et al. The cBio cancer genomics portal: an open platform for exploring multidimensional cancer genomics data. Cancer Discov 2012; 2(5): 401–404; doi 10.1158/2159-8290.CD-12-0095.

42. Gao J, Aksoy BA, Dogrusoz U, Dresdner G, Gross B, Sumer SO et al. Integrative analysis of complex cancer genomics and clinical profiles using the cBioPortal. Sci Signal 2013; 6(269): pl1. e-pub ahead of print 20130402; doi 10.1126/scisignal.2004088.

43. Brown DA, Di Cerbo V, Feldmann A, Ahn J, Ito S, Blackledge NP et al. The SET1 Complex Selects Actively Transcribed Target Genes via Multivalent Interaction with CpG Island Chromatin. Cell Rep 2017; 20(10): 2313–2327; doi 10.1016/j.celrep.2017.08.030.

44. Jette NR, Radhamani S, Arthur G, Ye R, Goutam S, Bolyos A et al. Combined poly-ADP ribose polymerase and ataxia-telangiectasia mutated/Rad3-related inhibition targets ataxia-telangiectasia mutated-deficient lung cancer cells. Br J Cancer 2019; 121(7): 600–610; e-pub ahead of print 2019/09/05; doi 10.1038/s41416-019-0565-8.

45. Guillemette S, Serra RW, Peng M, Hayes JA, Konstantinopoulos PA, Green MR et al. Resistance to therapy in BRCA2 mutant cells due to loss of the nucleosome remodeling factor CHD4. Genes Dev 2015; 29(5): 489–494; e-pub ahead of print 2015/03/05; doi 10.1101/gad.256214.114.

46. Hong R, Ma F, Zhang W, Yu X, Li Q, Luo Y et al. 53BP1 depletion causes PARP inhibitor resistance in ATM-deficient breast cancer cells. BMC Cancer 2016; 16(1): 725; e-pub ahead of print 20160909; doi 10.1186/s12885-016-2754-7.

47. Murai J, Huang SY, Das BB, Renaud A, Zhang Y, Doroshow JH et al. Trapping of PARP1 and PARP2 by Clinical PARP Inhibitors. Cancer Res 2012; 72(21): 5588–5599; doi 10.1158/0008-5472.CAN-12-2753.

48. Kranz A, Anastassiadis K. The role of SETD1A and SETD1B in development and disease. Biochim Biophys Acta Gene Regul Mech 2020; 1863(8): 194578; e-pub ahead of print 20200508; doi 10.1016/j.bbagrm.2020.194578.

49. Wyatt HD, West SC. Holliday junction resolvases. Cold Spring Harb Perspect Biol 2014; 6(9): a023192; e-pub ahead of print 20140902; doi 10.1101/cshperspect.a023192.

50. Abraham J, Lemmers B, Hande MP, Moynahan ME, Chahwan C, Ciccia A et al. Eme1 is involved in DNA damage processing and maintenance of genomic stability in mammalian cells. EMBO J 2003; 22(22): 6137–6147; doi 10.1093/emboj/cdg580.

51. Wyatt HD, Sarbajna S, Matos J, West SC. Coordinated actions of SLX1-SLX4 and MUS81-EME1 for Holliday junction resolution in human cells. Mol Cell 2013; 52(2): 234–247; e-pub ahead of print 20130926; doi 10.1016/j.molcel.2013.08.035.

52. Bhin J, Paes Dias M, Gogola E, Rolfs F, Piersma SR, de Bruijn R et al. Multi-omics analysis reveals distinct non-reversion mechanisms of PARPi resistance in BRCA1-versus BRCA2-deficient mammary tumors. Cell Rep 2023; 42(5): 112538; e-pub ahead of print 20230519; doi 10.1016/j.celrep.2023.112538.

53. Gogola E, Duarte AA, de Ruiter JR, Wiegant WW, Schmid JA, de Bruijn R et al. Selective Loss of PARG Restores PARylation and Counteracts PARP Inhibitor-Mediated Synthetic Lethality. Cancer Cell 2018; 33(6): 1078–1093 e1012; doi 10.1016/j.ccell.2018.05.008.

54. Chan KH, Rutazanaa D, Wray C, Thosani N, Yang V, Cen P. Promising Response of Olaparib in Patient With Germline ATM-Mutated Metastatic Gastric Cancer. J Investig Med High Impact Case Rep 2024; 12: 23247096241240176; doi 10.1177/23247096241240176.

55. Joris S, Denys H, Collignon J, Rasschaert M, T’Kint de Roodenbeke D, Duhoux FP et al. Efficacy of olaparib in advanced cancers with germline or somatic mutations in BRCA1, BRCA2, CHEK2 and ATM, a Belgian Precision tumor-agnostic phase II study. ESMO Open 2023; 8(6): 102041; e-pub ahead of print 20231016; doi 10.1016/j.esmoop.2023.102041.

56. Phan Z, Ford CE, Caldon CE. DNA repair biomarkers to guide usage of combined PARP inhibitors and chemotherapy: A meta-analysis and systematic review. Pharmacol Res 2023; 196: 106927; e-pub ahead of print 20230917; doi 10.1016/j.phrs.2023.106927.

57. Su CT, Nizialek E, Berchuck JE, Vlachostergios PJ, Ashkar R, Sokolova A et al. Differential responses to taxanes and PARP inhibitors in ATM-versus BRCA2-mutated metastatic castrate-resistant prostate cancer. Prostate 2023; 83(3): 227–236; e-pub ahead of print 20221116; doi 10.1002/pros.24454.

58. Gilmer TM, Lai CH, Guo K, Deland K, Ashcraft KA, Stewart AE et al. A novel dual ATM/DNA-PK inhibitor, XRD-0394, potently radiosensitizes and potentiates PARP and topoisomerase I inhibitors. Mol Cancer Ther 2024; e-pub ahead of print 20240408; doi 10.1158/1535-7163.MCT-23-0890.

59. Zhang A, Zhang L, Xie X, Liu D. Inhibition of ATM with KU-55933 Sensitizes Endometrial Cancer Cell Lines to Olaparib. Onco Targets Ther 2023; 16: 1061–1071; e-pub ahead of print 20231219; doi 10.2147/OTT.S426923.

60. Zhou Y, Borcsok J, Adib E, Kamran SC, Neil AJ, Stawiski K et al. ATM deficiency confers specific therapeutic vulnerabilities in bladder cancer. Sci Adv 2023; 9(47): eadg2263; e-pub ahead of print 20231122; doi 10.1126/sciadv.adg2263.

61. D’Ambrosio C, Erriquez J, Capellero S, Cignetto S, Alvaro M, Ciamporcero E et al. Cancer Cells Haploinsufficient for ATM Are Sensitized to PARP Inhibitors by MET Inhibition. Int J Mol Sci 2022; 23(10); e-pub ahead of print 20220521; doi 10.3390/ijms23105770.

62. Balmus G, Pilger D, Coates J, Demir M, Sczaniecka-Clift M, Barros AC et al. ATM orchestrates the DNA-damage response to counter toxic non-homologous end-joining at broken replication forks. Nat Commun 2019; 10(1): 87; e-pub ahead of print 20190108; doi 10.1038/s41467-018-07729-2.

63. Wyatt HD, Laister RC, Martin SR, Arrowsmith CH, West SC. The SMX DNA Repair Tri-nuclease. Mol Cell 2017; 65(5): 848–860 e811; doi 10.1016/j.molcel.2017.01.031.

64. Rondinelli B, Gogola E, Yucel H, Duarte AA, van de Ven M, van der Sluijs R et al. EZH2 promotes degradation of stalled replication forks by recruiting MUS81 through histone H3 trimethylation. Nat Cell Biol 2017; 19(11): 1371–1378; e-pub ahead of print 20171016; doi 10.1038/ncb3626.

65. Pepe A, West SC. Substrate specificity of the MUS81-EME2 structure selective endonuclease. Nucleic Acids Res 2014; 42(6): 3833–3845; e-pub ahead of print 20131225; doi 10.1093/nar/gkt1333.

66. Ray Chaudhuri A, Callen E, Ding X, Gogola E, Duarte AA, Lee JE et al. Replication fork stability confers chemoresistance in BRCA-deficient cells. Nature 2016; 535(7612): 382–387; e-pub ahead of print 2016/07/23; doi 10.1038/nature18325.

67. Panzarino NJ, Krais JJ, Cong K, Peng M, Mosqueda M, Nayak SU et al. Replication Gaps Underlie BRCA Deficiency and Therapy Response. Cancer Res 2021; 81(5): 1388–1397; e-pub ahead of print 2020/11/14; doi 10.1158/0008-5472.CAN-20-1602.

