## Supplemental Figures for "SETD1A-dependent EME1 transcription drives PARPi sensitivity in HR deficient tumour cells"

#### Supporting Information

**Figure S1 – Verification of SETD1A, ATM and BRCA1 depletion and the consequences for DNA damage signalling.**

**(A-B)** HeLa cells from Fig 1A (A) and Fig 1D (B) were irradiated with 10 Gy of ionising radiation, left for 1 hour, and whole cells extracts analysed by immunoblotting with the denoted antibodies. **(C-D)** Whole cell extracts of HeLa cells from Fig 2F (C) and 2G (D) were analysed by immunoblotting with the indicated antibodies.

**Figure S2 – Loss of SETD1A in ATM-deficient cells does not impact on PARP1 trapping by Olaparib.**

**(A)** Inducible HeLa Kyoto iCas9 cells expressing SETD1A gRNA were treated ± 1 μg/ml doxycycline, incubated for 48 hours, and then treated with 10 μM Olaparib ± 1 μM AZD0156 for a further 24 hours. Whole cell, nuclear soluble and chromatin bound fractions were prepared and assessed by immunoblotting using the denoted antibodies. **(B)** Band intensities were quantified using ImageJ. Data represent mean ± SEM from 4 independent replicates.

**Figure S3 – Validation of HeLa Kyoto iCas9 cells expressing SETD1A gRNA.**

(A-D) Inducible HeLa Kyoto iCas9 cells expressing SETD1A gRNA were treated ± 1 μg/ml doxycycline for 48 h. Cells were seeded onto coverslips and exposed to 3 Gy ionising radiation (IR), left for 8 h, and immunostained with antibodies against CENPF and BRCA1 or RIF1. Foci were enumerated in CENPF-negative G1 cells (A and B). Representative images are shown in C and D. **(E-H)** Whole cell extracts of cells from Fig 5A were analysed by immunobotting using the denoted antibodies (E). Band intensities were quantified using ImageJ (F-H). Data represent mean ± SEM from 3 independent replicates. * = $P < 0.05$, ** = $P < 0.01$, ***= $P < 0.0005$, ****= $P < 0.0001$ as determined by a one-way ANOVA with post-hoc Tukey test for multiple comparisons.

**Figure S4 – Validation of RNA-Seq data and qPCR analysis of differentially expressed genes.**

**(A)** Log2 counts per million (CPM) for the indicated differentially expressed transcripts identified from RNA-Seq analysis in Fig 5A. (B) mRNA was isolated from cells from Fig 5A, and mRNA expression levels of the denoted transcripts determined by qPCR. Data represent the mean ± SEM from five independent biological repeats. * = $P < 0.05$, ** = $P < 0.01$, ***= $P < 0.0005$, ****= $P < 0.0001$ as determined by a two-tailed unpaired Students t-test.

**Fig S5 – Localisation of SETD1A to the TSS of differentially expressed genes.**

Chromatin immunoprecipitation profiles of murine Setd1A at the TSS of genes identified as differentially regulated by loss of SETD1A from Fig 5A-C. Data is from [43].

**Fig S6 – Expression of SETD1A-dependent transcripts in lung, breast and ovarian cancer patient cohorts.**

**(A-C)** Log2 mRNA expression of genes identified in Fig 5A-C in patients diagnosed with (A) lung adenocarcinoma, (B) breast carcinoma, or (C) ovarian carcinoma patients triaged by SETD1A expression. Low= homozygous SETD1A deletion or mRNA expression <=-2 SD below mean; high= SETD1A mRNA expression >2 SD above mean. Datasets were obtained from CBioPortal [41,42]. ** = P

<0.01, ***= P < 0.0005, ****= P < 0.0001 as determined by a one-way ANOVA with post-hoc Tukey test for multiple comparisons.

**Fig S7 – Impact of SETD1A or EME1 deficiency on overall survival of lung, breast and ovarian cancer patients.**

**(A-B)** Kaplan-Meir curves denoting overall survival of BRCA1/2 wild-type breast cancer patients (A) or ATM-proficient lung adenocarcinoma patients (B) stratified by triaged by SETD1A expression as in Fig 5G. **(C-F)** Kaplan-Meir curves denoting overall survival of the denoted cancer subtypes (as in Fig 5G) stratified by triaged by EME1 expression. Low= homozygous EME1 deletion or mRNA expression <=-2 SD below mean; high= SETD1A mRNA expression >2 SD above mean. Datasets were obtained from CBioPortal [41,42]. * = P <0.05 as determined by a Log-rank (Mantel-Cox) test.

**Figure S1**

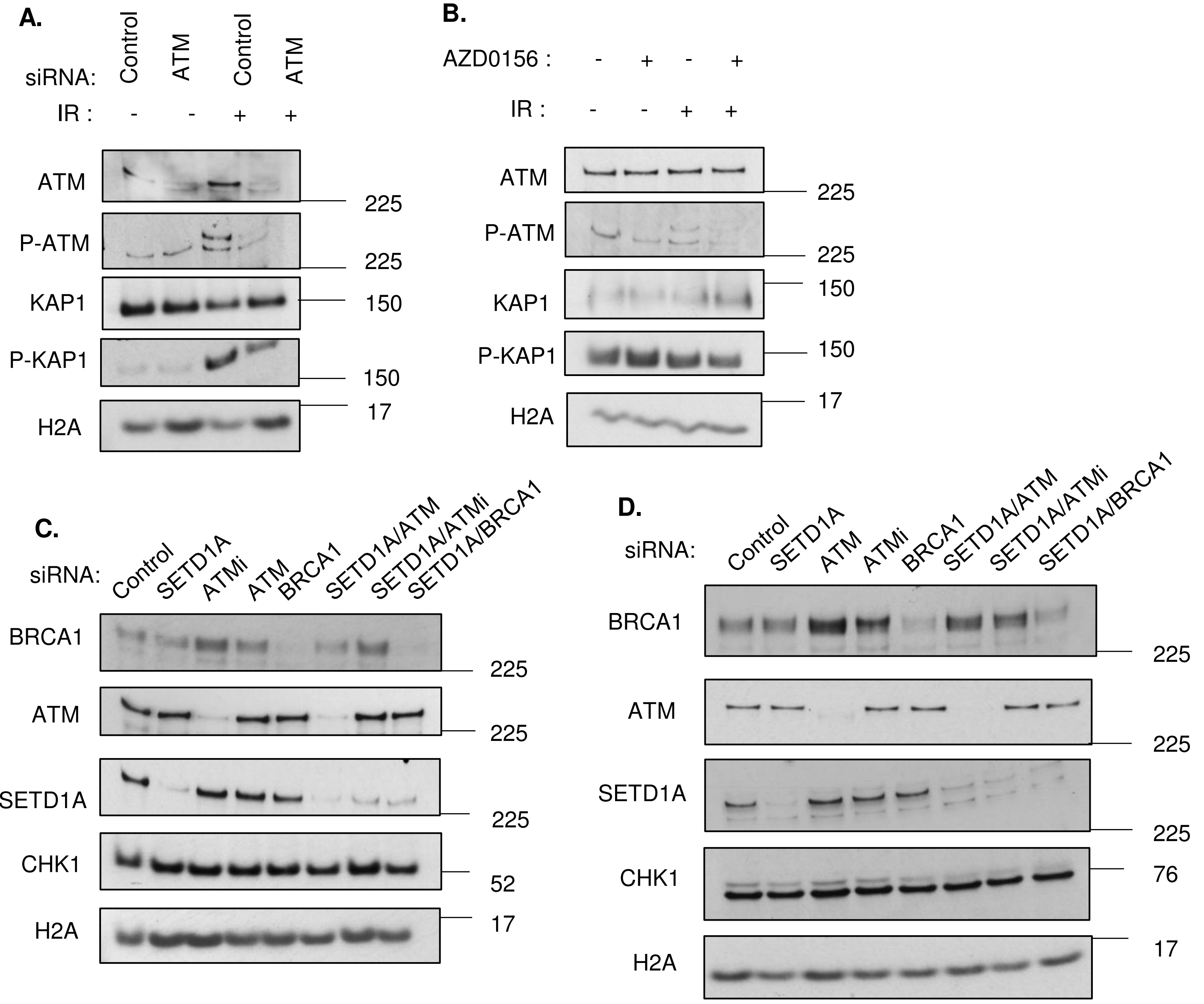

A.
siRNA:
Control
ATM
Control
ATM
IR :
- - + +
ATM
P-ATM
KAP1
P-KAP1
H2A
225
225
150
150
17
B.
AZD0156 :
- + - +
IR :
- - + +
ATM
P-ATM
KAP1
P-KAP1
H2A
225
225
150
150
17
C.
siRNA:
Control
SETD1A
ATMi
ATM
BRCA1
SETD1A/ATM
SETD1A/ATMi
SETD1A/BRCA1
BRCA1
ATM
SETD1A
CHK1
H2A
225
225
225
52
17
D.
siRNA:
Control
SETD1A
ATM
ATMi
BRCA1
SETD1A/ATM
SETD1A/ATMi
SETD1A/BRCA1
BRCA1
ATM
SETD1A
CHK1
H2A
225
225
225
76
17

### Figure S2

A.

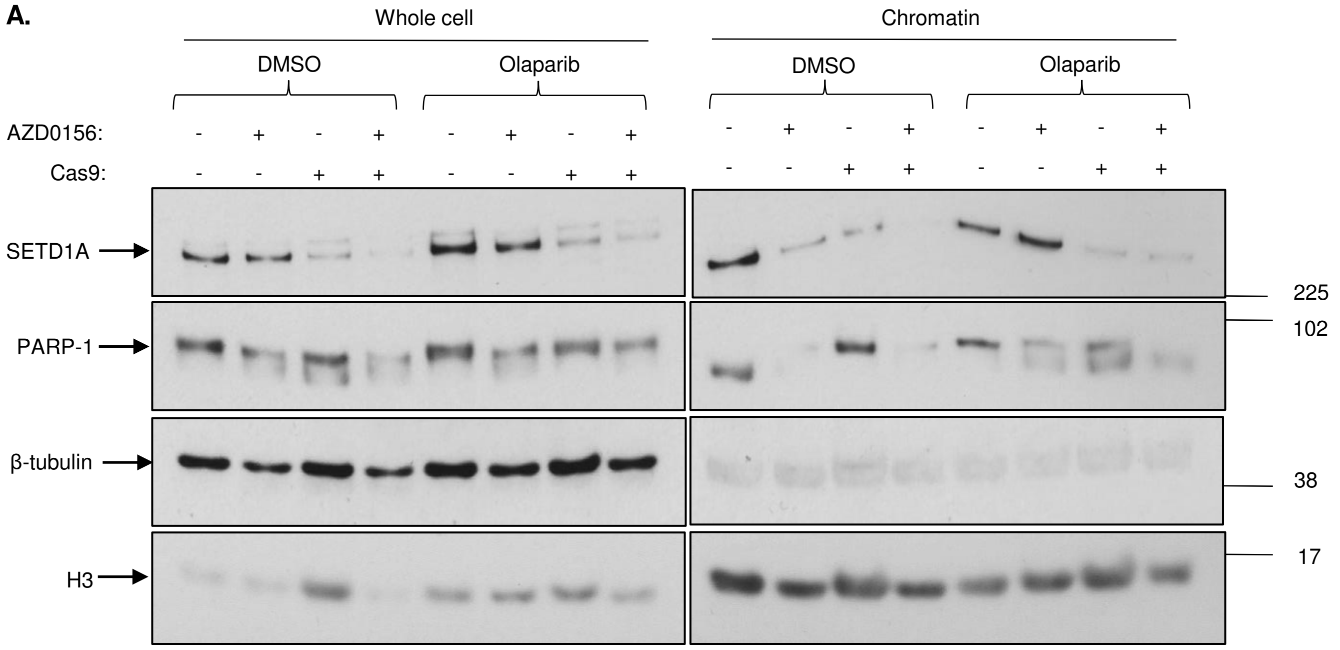
Whole cell
Chromatin
DMSO
Olaparib
DMSO
Olaparib
AZD0156:
Cas9:
SETD1A
PARP-1
β-tubulin
H3
225
102
38
17

B.

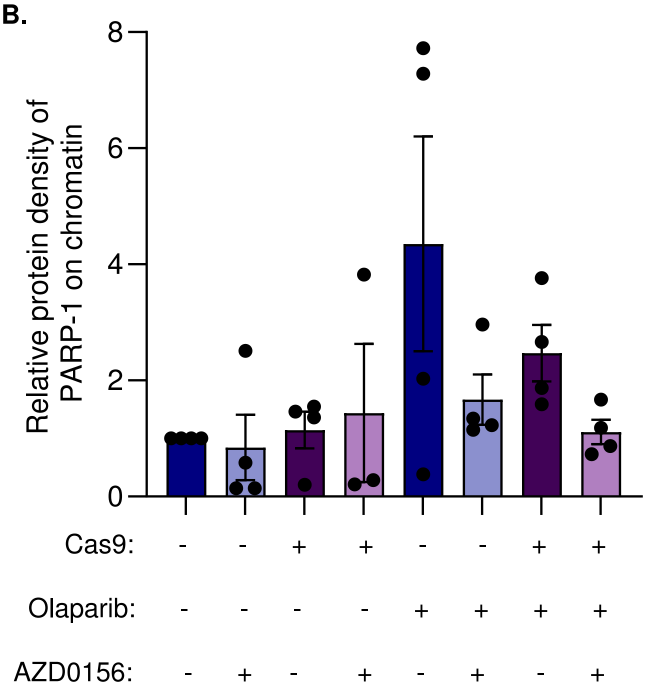
Relative protein density of PARP-1 on chromatin
8
6
4
2
0
Cas9:
Olaparib:
AZD0156:

#### Figure S3

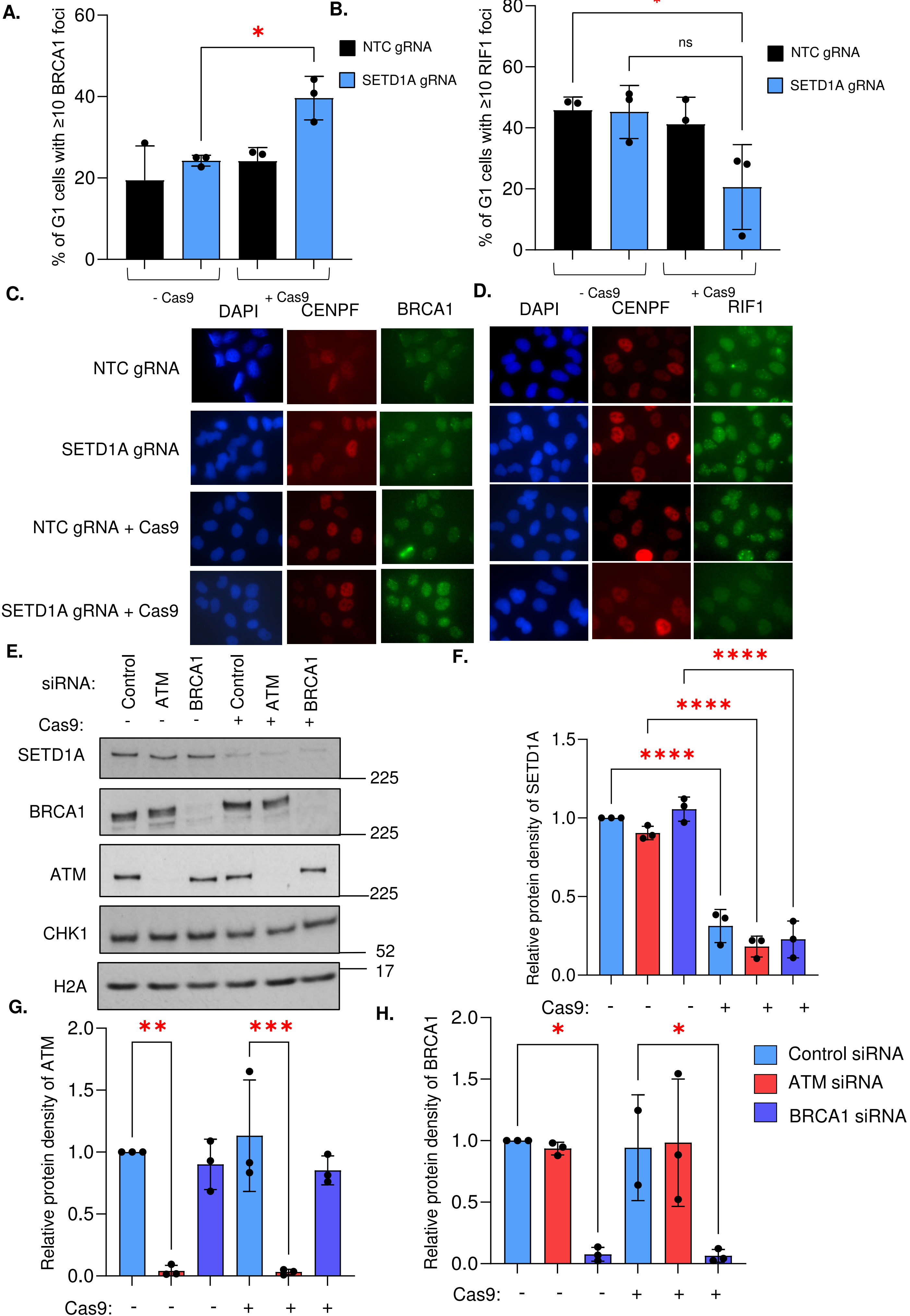

A.
% of G1 cells with ≥10 BRCA1 foci
60
40
20
0
*
- Cas9
+ Cas9
B.
NTC gRNA
SETD1A gRNA
% of G1 cells with ≥10 RIF1 foci
80
60
40
20
0
*
ns
- Cas9
+ Cas9
C.
DAPI
CENPF
BRCA1
NTC gRNA
SETD1A gRNA
NTC gRNA + Cas9
SETD1A gRNA + Cas9
D.
DAPI
CENPF
RIF1
E.
siRNA: Control ATM BRCA1 Control ATM BRCA1
Cas9: - - - + + +
SETD1A
BRCA1
ATM
CHK1
H2A
225
225
225
52
17
F.
Relative protein density of SETD1A
1.5
1.0
0.5
0.0
****
****
****
Cas9: - - - + + +
G.
Relative protein density of ATM
2.0
1.5
1.0
0.5
0.0
**
***
Cas9: - - - + + +
H.
Relative protein density of BRCA1
2.0
1.5
1.0
0.5
0.0
*
*
Cas9: - - - + + +
Control siRNA
ATM siRNA
BRCA1 siRNA

### Figure S4

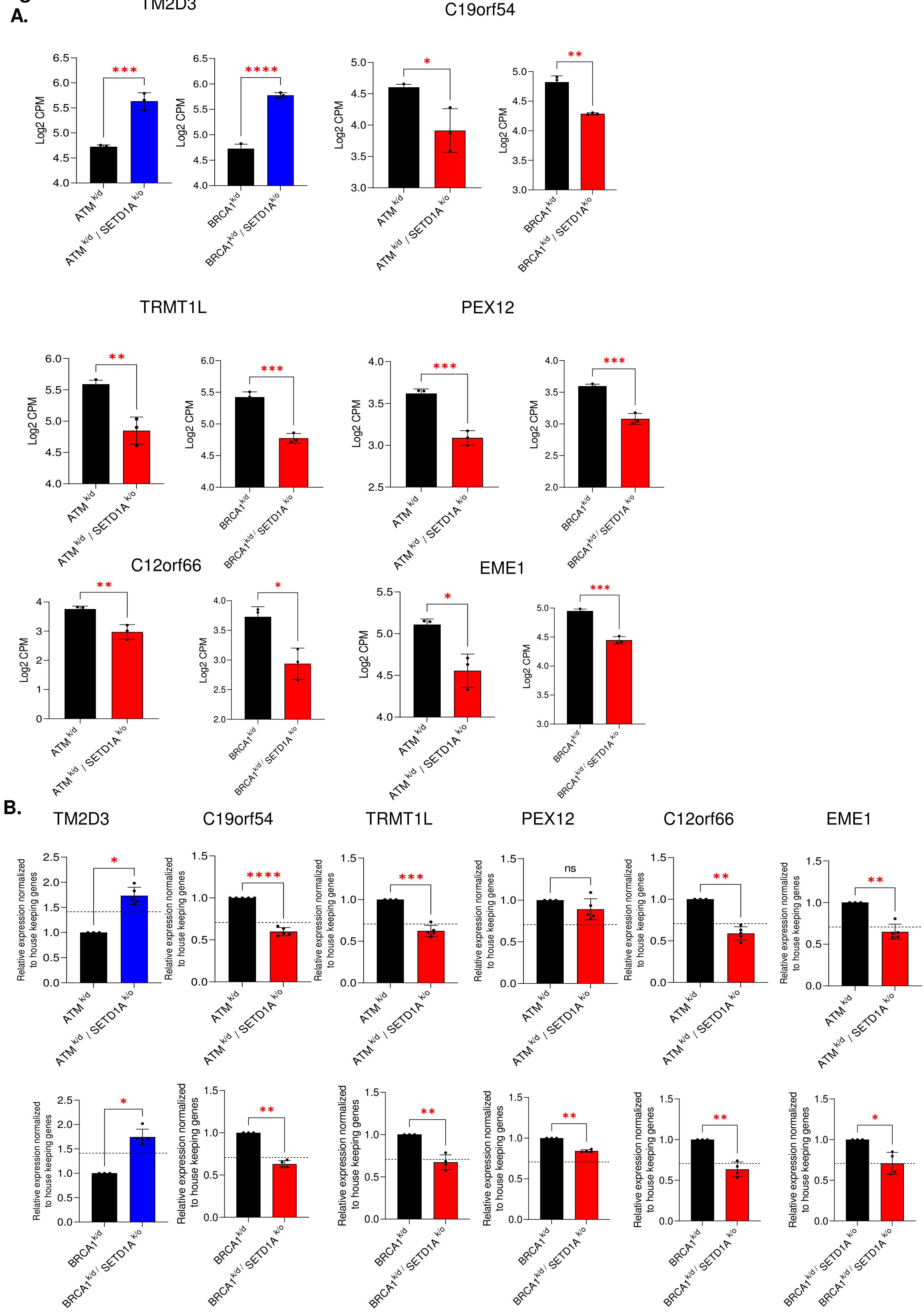

A.
TM2D3
C19orf54
TRMT1L
PEX12
C12orf66
EME1
Log2 CPM
ATM k/d
ATM k/d / SETD1A k/o
BRCA1 k/d
BRCA1 k/d / SETD1A k/o
B.
TM2D3
C19orf54
TRMT1L
PEX12
C12orf66
EME1
Relative expression normalized to house keeping genes
ns

### Figure S5

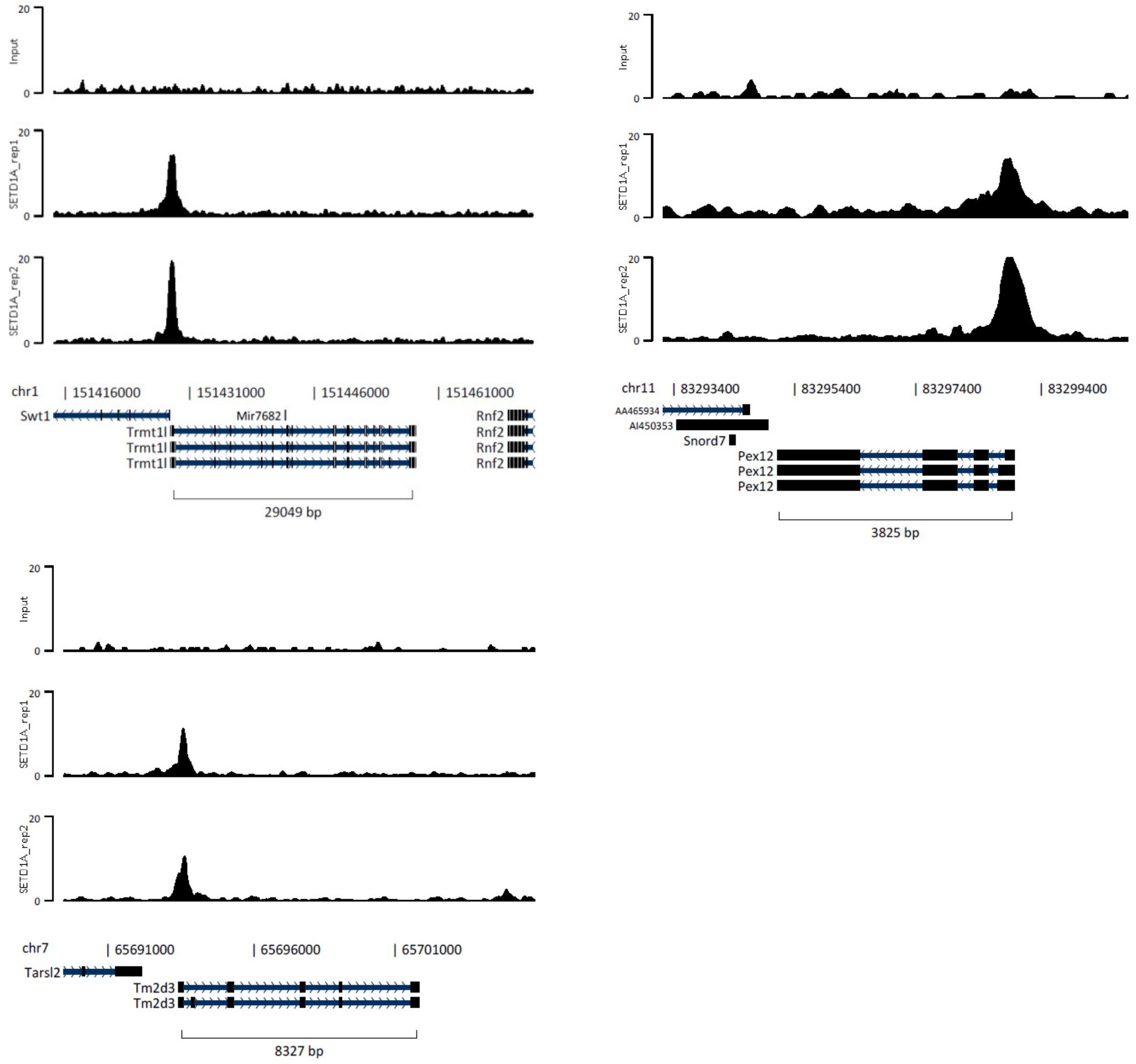

Input
SETD1A_rep1
SETD1A_rep2
20
0
chr1
151416000
151431000
151446000
151461000
Swt1
Mir7682
Trmt1l
Rnf2
29049 bp
chr11
83293400
83295400
83297400
83299400
AA465934
AI450353
Snord7
Pex12
3825 bp
chr7
65691000
65696000
65701000
Tarsl2
Tm2d3
8327 bp

#### Figure S6

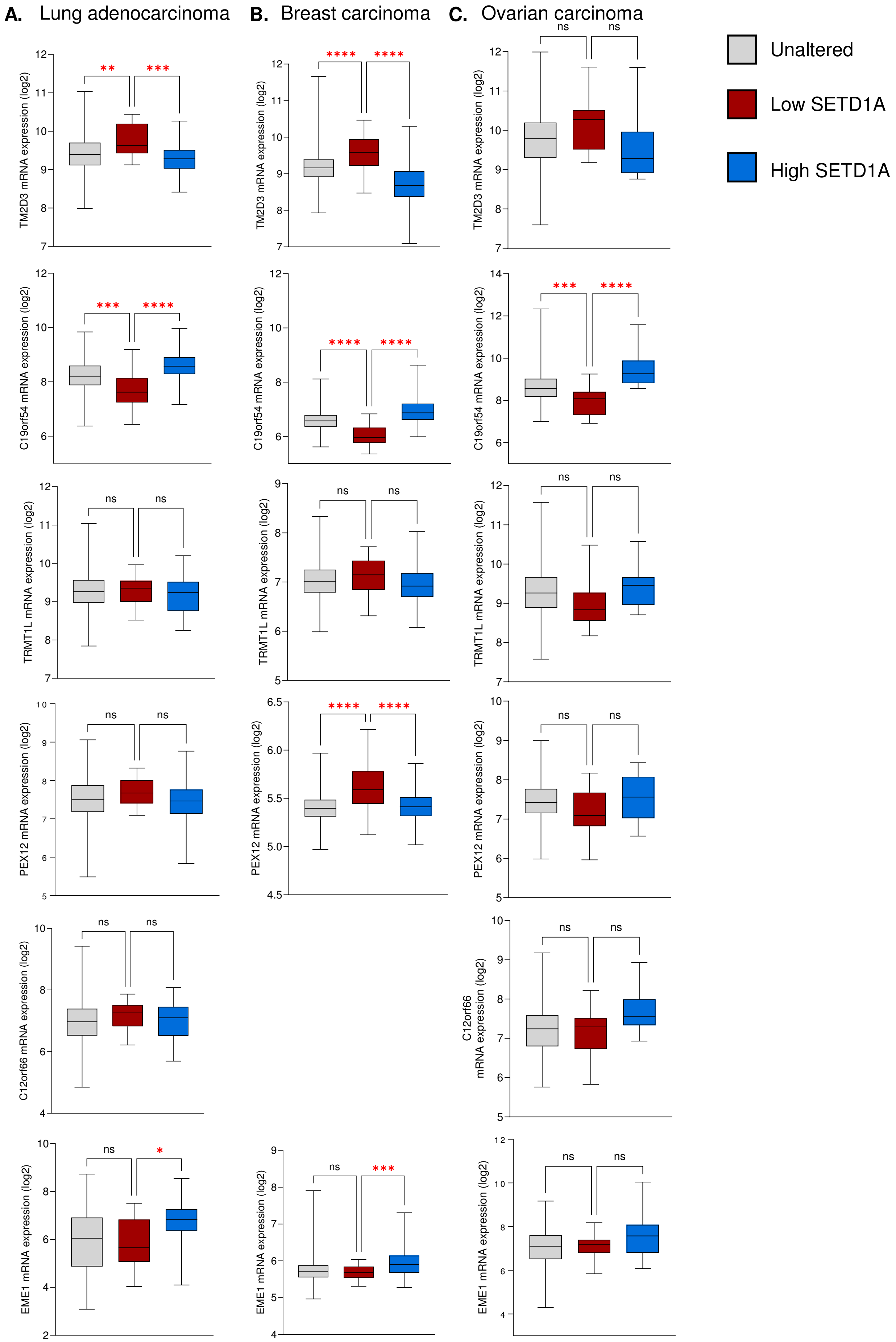
A. Lung adenocarcinoma
B. Breast carcinoma
C. Ovarian carcinoma
Unaltered
Low SETD1A
High SETD1A
TM2D3 mRNA expression (log2)
C19orf54 mRNA expression (log2)
TRMT1L mRNA expression (log2)
PEX12 mRNA expression (log2)
C12orf66 mRNA expression (log2)
EME1 mRNA expression (log2)
ns
*
**
***
****

### Figure S7

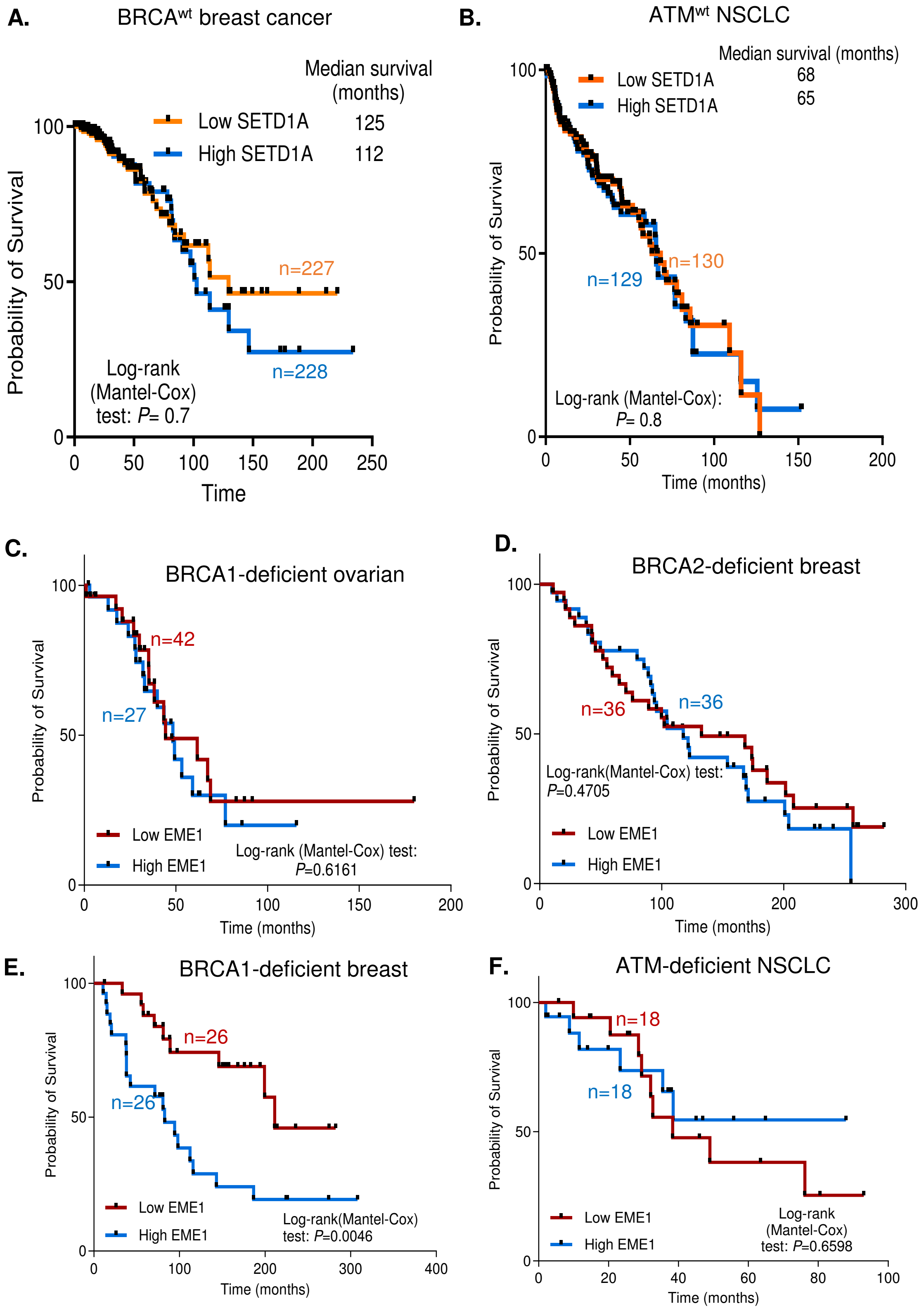
A.
BRCA$^{wt}$ breast cancer
Median survival (months)
Low SETD1A 125
High SETD1A 112
n=227
n=228
Log-rank (Mantel-Cox) test: P= 0.7
Probability of Survival
Time
100
50
0
0
50
100
150
200
250
B.
ATM$^{wt}$ NSCLC
Median survival (months)
Low SETD1A 68
High SETD1A 65
n=130
n=129
Log-rank (Mantel-Cox): P= 0.8
Probability of Survival
Time (months)
100
50
0
0
50
100
150
200
C.
BRCA1-deficient ovarian
n=42
n=27
Low EME1
High EME1
Log-rank (Mantel-Cox) test: P=0.6161
Probability of Survival
Time (months)
100
50
0
0
50
100
150
200
D.
BRCA2-deficient breast
n=36
n=36
Log-rank(Mantel-Cox) test: P=0.4705
Low EME1
High EME1
Probability of Survival
Time (months)
100
50
0
0
100
200
300
E.
BRCA1-deficient breast
n=26
n=26
Low EME1
High EME1
Log-rank(Mantel-Cox) test: P=0.0046
Probability of Survival
Time (months)
100
50
0
0
100
200
300
400
F.
ATM-deficient NSCLC
n=18
n=18
Low EME1
High EME1
Log-rank (Mantel-Cox) test: P=0.6598
Probability of Survival
Time (months)
100
50
0
0
20
40
60
80
100
